# Focal Infrared Neural Stimulation Propagates Dynamical Transformations in Auditory Cortex

**DOI:** 10.1101/2025.03.12.642906

**Authors:** Brandon S Coventry, Cuong P Luu, Edward L Bartlett

**Affiliations:** Weldon School of Biomedical Engineering, Purdue University, West Lafayette, IN 47907 USA; Center for Implantable Devices, Purdue University, West Lafayette, IN 47907 USA; Institute for Integrative Neuroscience, Purdue University, West Lafayette, IN 47907 USA; School of Medicine and Public Health, University of Wisconsin-Madison, Madison, WI 53907 USA; Department of Biological Sciences, Purdue University, West Lafayette, IN 47907 USA

**Keywords:** Infrared Neural Stimulation, Thalamocortical Circuits, Deep Brain Stimulation, Cochlear Implant, Cortex, Chaos, Thalamus

## Abstract

**Significance:** Infrared neural stimulation (INS) has emerged as a potent neuromodulation technology, offering safe and focal stimulation with superior spatial recruitment profiles compared to conventional electrical methods. However, the neural dynamics induced by INS stimulation remain poorly understood. Elucidating these dynamics will help develop new INS stimulation paradigms and advance its clinical application.

**Aim:** In this study, we assessed the local network dynamics of INS entrainment in the auditory thalamocortical circuit using the chronically implanted rat model; our approach focused on measuring INS energy-based local field potential (LFP) recruitment induced by focal thalamocortical stimulation. We further characterized linear and nonlinear oscillatory LFP activity in response to single-pulse and periodic INS and performed spectral decomposition to uncover specific LFP band entrainment to INS. Finally, we examined spike-field transformations across the thalamocortical synapse using spike-LFP coherence coupling.

**Results:** We found that INS significantly increases LFP amplitude as a log-linear function of INS energy per pulse, primarily entraining to LFP *β* and *γ* bands with synchrony extending to 200 Hz in some cases. A subset of neurons demonstrated nonlinear, chaotic oscillations linked to information transfer across cortical circuits. Finally, we utilized spike-field coherences to correlate spike coupling to LFP frequency band activity and suggest an energy-dependent model of network activation resulting from INS stimulation.

**Conclusions:** We show that INS reliably drives robust network activity and can potently modulate cortical field potentials across a wide range of frequencies in a stimulus parameter-dependent manner. Based on these results, we propose design principles for developing full coverage, all-optical thalamocortical auditory neuroprostheses.

## 1 Introduction

Infrared neural stimulation (INS) is a potent optical stimulation modality, utilizing coherent infrared light^1,2^ to drive excitatory^3–5^ and inhibitory^6^ responses in nerves^7–10^, neurons^3,11,12^, and other excitable cells^13^. Compared to conventional electrical stimulation, INS demonstrates an advantageous spatial recruitment profile with limited spread of off-target activation^3,14,15^. Furthermore, INS safety^16,17^ and ablation thresholds^18,19^ have been well characterized in preclinical animal models and preliminary human studies. Advancements in implantable fiber optics satisfying the technical demands of therapeutic laser stimulation^20^ have made INS an attractive stimulation paradigm for clinical neural modulation in cochlear implants^2,21^, deep brain stimulation^3,17^, and peripheral nerve stimulation^7,22–24^. However, while INS in the human cerebral cortex has shown focal and reliable responses^17^, limited mechanistic understanding of neural entrainment to INS activation has restricted further translation of INS neural modulation into therapeutic use. Our previous work has shown that thalamocortical INS stimulation evokes short-latency spiking in auditory cortex neurons^3^. Other studies using patch clamp have shown robust inward depolarizing currents in response to infrared stimuli^4,25,26^, with some evidence of synaptic modulation at the level of the single neuron^25^. Studies aimed at elucidating the primary driver of neural activation have pointed to thermal gradients^27–29^, modulation of cellular lipid bilayers^13,30,31^, induction of intracellular calcium cycling^32,33^, and direct modulation of voltage gated channels^34^ as potential primary drivers of INS neural recruitment. Regardless of the mechanism of excitation, how INS activates brain networks that generates or resets cortical rhythms remains largely unknown. The vast majority of studies of cortical recruitment with INS utilize intracortical INS rather than the natural thalamic inputs to the cortex^11,17,35–39^. While these studies have been foundational in the understanding of INS recruitment of large-scale cortical circuits, studies which utilize thalamic and subthalamic inputs traditionally employed for clinical neuromodulation are necessary to advance knowledge of INS network entrainment and to put INS studies in alignment with current clinical neuromodulation therapies. Such knowledge will facilitate the optimization of INS parameters and design as well as further identification of therapeutic neural targets for clinical use, given that cortical rhythms are critical for the coordination of cortical activities^40–43^. Validation and characterization of cross-network INS entrainment can also position INS as a powerful tool for network neuroscience studies^3,44^.

Thalamocortical transformations are critical to the function of higher order neural circuits, performing complex coding to facilitate feedforward fine-tuned encoding of sensory and perceptual input^45–48^ with robust intracortical and corticofugal feedback projections allowing for both local and widespread regulation and plasticity^49–53^. Cortex wide processing is often assessed through measurement of local field potentials (LFPs), which function as a readout of grouped electric field activity^54^ and closely related transmembrane currents^55–57^. Analysis of band-limited oscillatory LFP activity in *θ* (4-8Hz), *α* (8-13Hz), *β*(13-30Hz), and *γ* (30-200Hz) can be utilized to assess thalamocortical encoding mechanisms, information transmission, and gain modulation^58^ as well as serving as discriminative biomarkers for disease state^59^. Modulation of thalamocortical network function has also been implicated as an important biomarker in therapeutic deep brain stimulation^60–64^, exemplified by effective DBS stimulation showing rapid reduction in *β* band activity and transient increases in high *γ* band activity in Parkinson’s disease^65–67^. Therefore, to realize the clinical and basic science potential of INS, a fuller understanding of the INS parameter-dependent induced network transformations is necessary.

In this study, we assess thalamocortical transformation and encoding of INS stimulation through chronic local field potential recordings in the rat. We quantify LFP INS dose-response characteristics while showing entrainment of INS primarily within *β* and *γ* LFP frequency bands. We also utilized LFP and single unit correlations to quantify dynamic transformations in the auditory thalamocortical circuit. To assess nonlinear dynamics of INS stimulation, we utilize tests for chaos dynamics to show that the auditory cortex contains neurons which exhibit chaotic neural dynamics that are directly modulated by INS stimulation of the thalamus. We further quantify single-unit spike contribution to LFP oscillatory activity through spike-field coherence measures, showing spike entrainment primarily through *β* and *γ* bands. Finally, owing to the observed neural encoding dynamics in this study, we provide guidance for use of INS as a thalamic auditory neuroprosthesis.

## 2 Materials and Methods

All experimental and surgical procedures and protocols were approved by the Institutional Animal Care and Use Committee (PACUC) of Purdue University (West Lafayette, IN, #120400631) and in accordance with the guidelines of the American Association for Laboratory Animal Science (AALAS) and the National Institutes of Health guidelines for animal research.

### 2.1 Surgical Procedures

Detailed INS optrode and electrode implant procedures can be found in the following protocol^68^. Briefly, adult Sprague Dawley rats weighing between 300-400g (Envigo, Indianapolis IN) were initially anesthetized with 5% isoflurane in *O*_2_ (1.2 mL/min flow rate) followed by a bolus intramuscular injection of a ketamine/dexmedetomidine cocktail (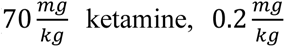 dexmedetomidine) to achieve a surgical plane of anesthesia. Rats were given an intramuscular injection of Buprenorphine 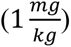 for analgesia at least 30 minutes prior to first incision. After induction, rats were transferred to a stereotaxic frame secured by hollow ear bars. Animal heart rate and pulse oximetry were monitored throughout the duration of the surgery and recovery, with anesthetic surgical plane assessed via lack of toe-pinch reflex every 15 minutes. An initial incision was made over midline and the periosteum removed via blunt dissection to reveal cranial sutures. Three stainless steel bone screws were placed in the skull to provide stability for implanted devices and headcap with a fourth titanium bone screw secured to the skull to serve as recording electrode ground and reference. A craniectomy was made above auditory thalamus (-6 anterior/posterior, - 3.5 medial/lateral, -6 dorsal/ventral) and a fiber optic optrode (Thor Labs, Newton, NJ, USA, 200*μm* fiber diameter, 0.22 NA) was slowly advanced and placed in the medial geniculate body. The optrode was sealed into place using UV-curable dental acrylic (Pentron, Wallingford, CT, USA). Next the right temporalis muscle was gently reflected off the skull and a 2x2 mm craniectomy was made over auditory cortex. Dura mater was gently resected using a 25G bent tip needle. A 2 mm × 2 mm 16 channel microwire array (TDT, Alachua, FL, USA, electrode spacing given in Fig. 1A) was slowly inserted perpendicular into auditory cortex. Broadband 80dB-SPL gaussian-noise stimuli were applied to the contralateral ear during electrode insertion to putatively confirm putative placement of recording arrays into primary afferent layer III/IV as evidenced by recordings of short latency, high amplitude multiunit activity entrained to auditory stimuli. One animal received 3 mm linear recording array (NeuroNexus A1-16, 200*μm* between contacts) in layer III/IV of auditory cortex. Devices were sealed into placed by the application of Qwik-Sil (World-Precision Instruments) to seal the craniectomy followed by application of UV-curable dental acrylic. A low profile headcap was completed to ensure device stability. After the completion of the surgery, animals were returned to their home cage and allowed to recover for at least 72 hours. Supplemental Buprenorphine analgesia was administered every 6-12 hours for a minimum of 72 hours. A total of 5 rats had functional longitudinal optrode and electrode implants and were included in the study.

**Fig. 1.**
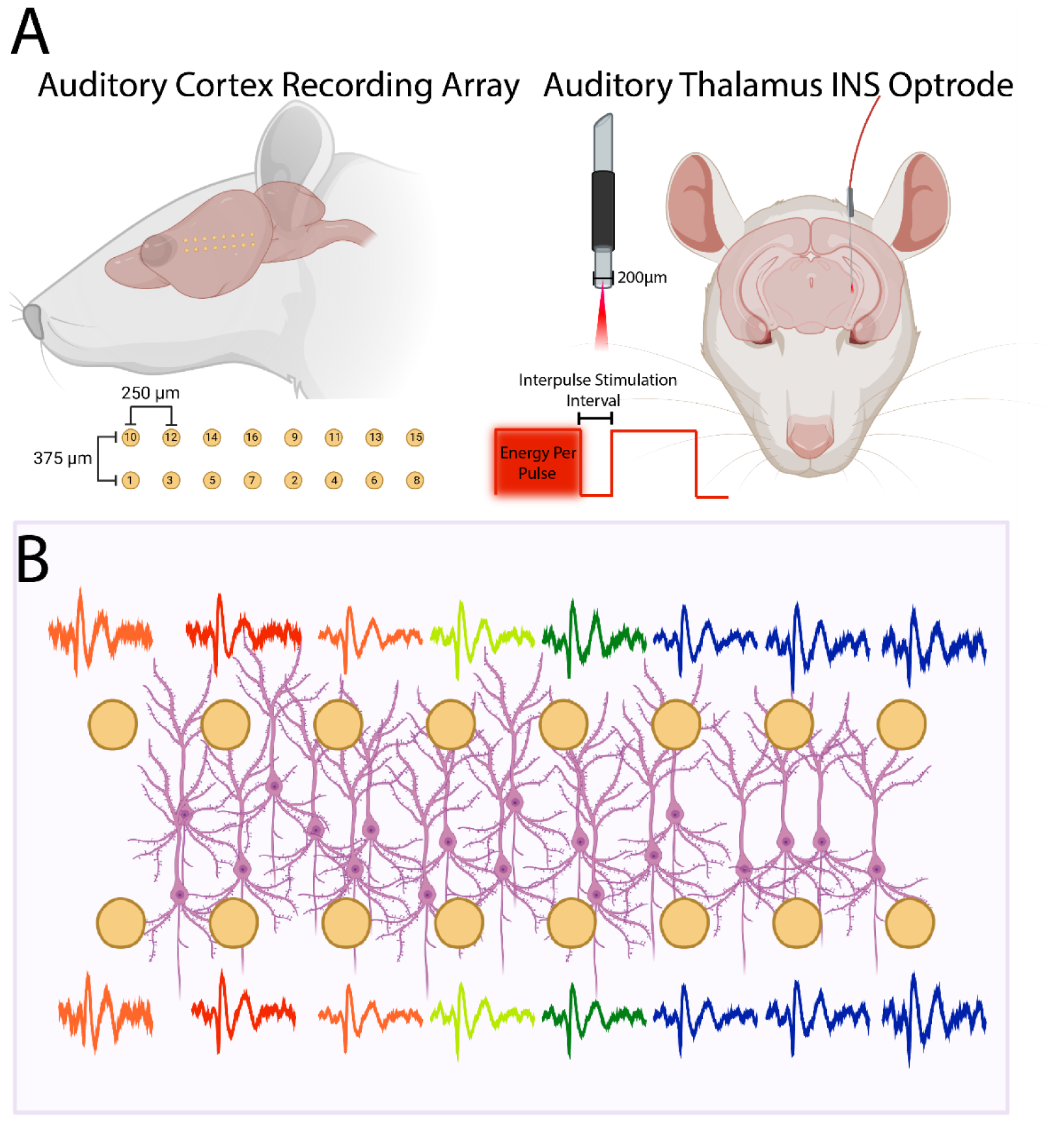
Schematic representation of stimulation and recording preparation. A. Rats were implanted with planar microwire arrays into layer III/IV of auditory cortex. Spacing of recording electrodes is schematized in this figure. Infrared optrodes were implanted into the medial geniculate body of auditory thalamus. Activation of thalamocortical loops was controlled by laser energy per pulse and interpulse stimulation intervals. B. Local field potential (LFP) recordings were made concurrently across all channels. Propagation of cortical waves was analyzed across all channels. Figure was crafted using tools from BioRender (www.biorender.com).

### 2.2 Infrared Stimulation

Optical stimulation was provided by INSight, our custom designed, open source optical neuromodulation platform^3,69^ using a 1907 nm semiconductor laser module(Akela Trio, Jamesburg, NJ, USA) fiber coupled to the implanted optrode(Thor Labs FG200LCC). Laser pulse train stimuli were controlled via analog outputs of a RX-7 stimulator base-station (Tucker-Davis Technologies, Alachua FL) with laser energies between 0-4 mJ constrained below ablation thresholds^19^ with interpulse stimulation intervals between 0.2 − 100ms. Optical stimulation was provided by INSight, our custom designed, open source optical neuromodulation platform^3,69^ using a 1907 nm semiconductor laser module(Akela Trio, Jamesburg, NJ, USA) fiber coupled to the implanted optrode(Thor Labs FG200LCC). Laser pulse train stimuli were controlled via analog outputs of a RX-7 stimulator base-station (Tucker-Davis Technologies, Alachua FL) with laser energies between 0-4 mJ constrained below ablation thresholds^19^ with interpulse stimulation intervals (ISI) between 0.2 − 100ms.

### 2.3 Electrophysiological recordings

LFP and multiunit activity was recorded across all channels of the implanted recording array (Fig 1B). All recordings were performed in a 9′ × 9′ electrically and acoustically isolated chamber (Industrial Acoustics Corporation, Naperville, IL, USA) with laser electronics placed outside of the chamber to prevent field interactions from large magnitude current pulses^70,71^. Rats were given intramuscular injections of Dexmedetomidine 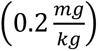 sedative^72–74^ to reduce movement artifact disruptions in neural recordings. Each recording trial was composed of a 200 ms prestimulus interval to facilitate spontaneous rate calculations followed by application INS with total trial lengths equal to 1s. INS laser energies were randomized to limit stimulus adaptation effects with 30-60 trials per INS energy. Signals from implanted recording arrays were amplified via a Medusa preamplification headstage (Tucker-Davis Technologies) and discretized with a sampling rate of 24.414kHz (TDT standard sampling rate) and recorded into a RZ-2 bioprocessor (Tucker-Davis Technologies). Real-time visualization of recorded signals was made using OpenEx software, with Spike channel bandpass filter cutoffs set to 0.3 − 5kHz and LFP channel bandpass filter cutoffs set to 3 − 300Hz.

### 2.4 Data Processing and Analyses

All data analysis was performed on a Windows-based computer with a Xeon 16 core processing and an Nvidia Titan X GPU for parallel and multithreaded processing. LFPs were down sampled to a sampling rate of 1526 Hz for analysis. Multiunit signals were exported and processed using custom-written programs in the Matlab programming environment (Mathworks, Natick MA). Multiunit activity was sorted into single units using wavelet-based superparamagnetic clustering methods in Wave-Clus^75^. Spikes were then derandomized and sorted into spike-time rasters and exported to Python for further analyses.

Time-domain INS dose response curves were quantified by first high-pass filtering raw LFPs to remove slow-baseline drifts (10 Hz filter cutoff, Chebyshev Type II high-pass filter). Evoked LFP onset amplitudes were quantified as the root-mean-squared (RMS) magnitude between stimulus onset through the first negative (N1)-second positive (P2) peaks. Dose-response characteristics were then quantified through Bayesian multilinear regressions (see 2.8 Statistical Inference).

For LFP frequency band analyses, raw mean LFPs were decomposed into *θ* (4-8Hz), *α* (8-13Hz), *β*(13-30Hz), low *γ* (30-80Hz), and high *γ* (80-200Hz) using the continuous fast wavelet transform^76^. CWTs were utilized over short-time Fourier transforms as CWTs have been found to provide better frequency resolution in non-stationary field potential recordings^77^. For frequency band analyses, the 10Hz high pass filter was not applied, with recorded signals only filtered by hardware high pass filters (cutoff = 3Hz). The total power in a frequency band was calculated as

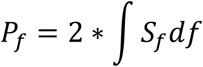

where 𝑆_𝑓_is the band-power spectral density from wavelet decomposition and the factor of 2 included to account for frequency folding inherent to time-frequency decomposition. Numerical integration was performed using Simpsons rule^78^. Stimulus-induced changes in frequency bands from baseline was quantified as:

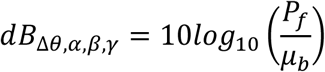

Where *μ*_𝑏_ is the mean power in the 200ms prestimulus window respectively.

### 2.5 Temporal Modulation Transfer Functions

To assess the sensitivity of INS-evoked LFPs to stimulus ISI, temporal modulation transfer functions (tMTF) were calculated. Fast-Fourier transforms (FFT) from raw LFPs were calculated using Bluestein’s method^79^ in Python. FFTs were calculated for baseline LFPs 200ms prior to stimulation onset and for stimulation windows consisting of the stimulation time + 100ms for longer lasting LFP responses. Spectral power at the stimulus ISI frequency was extracted from baseline and stimulation condtions with analysis completed for ISI frequencies < 763 Hz to ensure the Nyquist sampling criterion was met. Change in LFP power from baseline at the ISI frequency was calculated as:

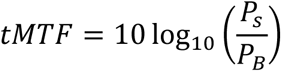

where 𝑃_𝑠_, 𝑃_𝐵_ are the LFP power at the ISI frequency for stimulation and baseline windows respectively. ISI modulation was considered significant if the tMTF had a value of ≥ 3 dB corresponding to a doubling in LFP-ISI power from baseline.

### 2.6 Estimation of Chaotic Cortical Dynamics

To estimate the criticality and chaoticity of LFP cortical dynamics, a modified 0-1 test was used^80–82^. The 0-1 test assesses the degree of chaotic dynamics in recorded time series of deterministic dynamical systems by mapping dynamics to a 2D space of the form:

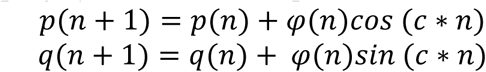

where φ(n) is the discretized LFP timeseries and c is a random value between 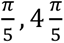. The range of c helps prevent overestimation due to mapping system resonance^81^. The dynamics of the system 𝑝(*n*), 𝑞(*n*) are such that regular, non-chaotic dynamics are bounded, and chaotic dynamics are unbounded and show movement similar to 2D Brownian motion process, and thus exhibit diffusive growth proportional to √*n*. ^81^. Examples of 𝑝(*n*), 𝑞(*n*) systems for non-chaotic and chaotic dynamics are shown in Fig S1. From the new 𝑝(*n*), 𝑞(*n*) system, the time-averaged mean squared displacement is calculated as:

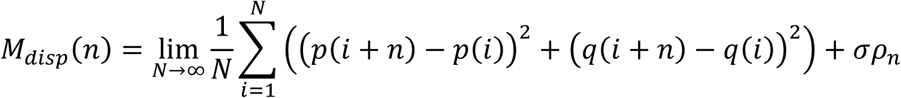

where 𝜌_*n*_ is a uniformly distributed random variable between 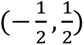 with 𝜎 the noise perturbation amplitude, set to 0.5 with N corresponding to the total number of samples contained in the time series. It can be shown that 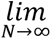 exists^81^. In practice, 𝑀_𝑑𝑖𝑠𝑝_ is calculated for *n* ≤ 𝑁_0_ where 𝑁_0_ ≪ 𝑁. A value of 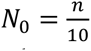 was chosen as it shows good performance on LFP timeseries^82,83^. The asymptotic growth rate of 𝑀_𝑑𝑖𝑠𝑝_ is then calculated as:

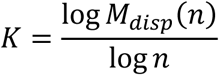

K thus describes a measure of “boundedness” of the system, scaled between (0,1), with 0 corresponding to bounded periodic dynamics and 1 corresponding to unbounded, chaotic dynamics. K values on the continuum between (0,1) thus represent degrees of unbounded growth. As periodic and aperiodic dynamics are sensitive to values of 𝑐, 𝐾 is computed for 100 iterations of the 0-1 test with c randomly drawn from a uniform distribution with support 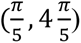. The median asymptotic growth rate coefficient is then calculated as:

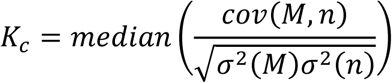

where 𝑀 = (𝑀_𝑑𝑖𝑠𝑝_(*n*)_𝑐1_, 𝑀_𝑑𝑖𝑠𝑝_(*n*)_𝑐2_, …, 𝑀_𝑑𝑖𝑠𝑝_(*n*)_𝑐100_) and 𝜎^2^ is the variance operator. Code for 0-1 testing was adapted and modified from the Chaos decision tree toolbox^82^.

Information content contained within LFPs was calculated using estimation of Shannon mutual information (MI). MI quantifies the reduction in uncertainty in neural responses (𝑅) given knowledge of a particular stimulus (𝑆_𝑥_) and thus quantifies thalamocortical encoding and separability of differing INS stimuli. Stimulus-Response 𝑝(𝑟|𝑠_𝑥_) and total response 𝑝(𝑟) probability density functions were estimated from total LFP responses using kernel density estimation with a total of 21 bins. Optimal bin counts were calculated using the Doane estimate^84^, which is robust for potentially non-normal distributions. MI was calculated using the methods of Borst and Theunissen^85^ as:

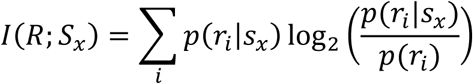

Bias in estimates of 𝐼(𝑅; 𝑆_𝑥_) due to imperfect knowledge of probability density functions was corrected using the methods of quadratic extrapolation^72,86,87^ as follows:

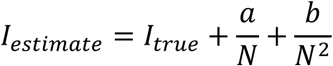

where N is the number of trials used in the MI estimate and 𝑎, 𝑏 are free parameters corresponding to stimulus-response dependence on sample size and is learned by least-squares regression from calculations of MI with sample sizes of 𝑁, 0.50𝑁, and 0.25𝑁. All information measures are reported after bias correction.

### 2.7 Spike-Field Coherence Measures

Spike-field coherence (SFC) measures were calculated to assess the band frequency in which spikes and LFPs are consistently entrained as a function of applied energy. Electrodes showing both LFP activity and graded single-unit firing rates in response to INS were included in this analysis. SFCs were calculated as:

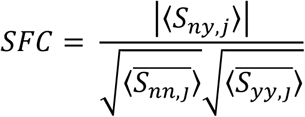

where 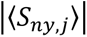 is the magnitude of the trial-averaged cross power spectral density between per- trial 𝑗𝑗 spike train *n* and LFP 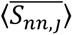 the trial-averaged auto power spectral density of spike train n across trial j, and 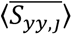 the trial-averaged auto power spectral density of the LFP y across trial j. SFCs were calculated using the Electrophysiology Analysis Toolbox^88^ synched in Python to SPyke across all applied INS energies.

### 2.8 Statistical Inference

Statistical inference was performed using Bayesian inference methods^89–92^. Bayesian methods allow for robust quantification of parameter effects and model uncertainty driven purely by observed data with robust model validation controls. Hierarchical Bayesian models are particularly suited for analysis of repeated measures designs and provides robust control of within and between subject variance^89^ such as this study. Significance of effect was assessed using observations of posterior distribution 95% highest density intervals and credible regions, in line with Bayesian convention^90–92^. Bayesian inference requires explicit declaration of prior distributions that quantify current state of knowledge. Prior distributions for each model are described below and are weakly informative and have minimal effect on posterior distributions owing to relative scarcity of data surrounding auditory cortex cortical oscillations in response to INS. Bayesian inference reporting follows the Bayesian analysis reporting guidelines outlined in ^93^ with posterior predictive checks of model fits to data and prior sensitivity checks reported in the supplementary material. Statistical significance is summarized in posterior maximum a priori (MAP) estimates of model parameters and posterior highest density intervals (HDIs) and credible regions in Bayesian inference convention.

Changes in LFP RMS magnitude and frequency band power as a function of applied energy were assessed via Bayesian Hierarchical Linear Regressions (BHLR). The hierarchical structure of BHLR allows for accommodation of within and between subject variance necessary for repeated measures designs. BHLR take the form of:

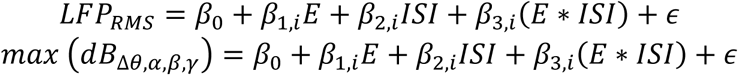

Where 𝐸 corresponds to energy per pulse, 𝐼𝑆𝐼 corresponding to interpulse stimulus interval, and (𝐸 ∗ 𝐼𝑆𝐼) corresponds to an energy per pulse and interpulse stimulus interval interactions. The *β*_0_and 𝜖 represent the regression intercept and model error respectively. Model comparison through leave-one out (LOO) criterion^94^ suggested that natural log transformation of response and independent variables provided regression models that best fit observed data (model comparisons show in supplementary information) in both regression modles. The *β*_𝑥,𝑖_terms correspond to regression slope coefficients for each factor for each electrode/animal group 𝑖. Full model structure is shown in Fig S2. Priors were set as broad, uninformative normal priors. Prior sensitivity analyses were performed to assess the quality of statistical model fit to data and are given in supplementary information. Regression parameters were considered significant if their 95% HDI and credible intervals, summarizing the range in which there is a 95% probability that the true estimate of the regression coefficient lies in the interval given evidence from observed data, does not contain 0 in line with Bayesian convention^90,93^. Results are presented as the maximum *a priori* estimate (95% credible interval) of each regression parameter.

LFP frequency band correlations were quantified as Pearson’s R correlation coefficient in the Scipy environment. Significance of correlation was assessed as a two-sided p-value, which if given the measured value R and a surrogate measure 𝑅, representing the correlation coefficient between shuffled input timeseries, is the probability of 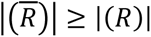. Significance level for correlation analysis was set to *α* = 0.05.

Spike rate, N1-P2 LFP RMS amplitude correlations were assessed using Bayesian linear regressions of the form:

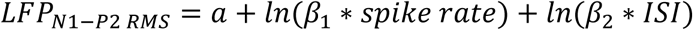

where 𝑎 is the regression intercept coefficient, spike rate corresponds to average spike rate within the stimulus interval plus a 50 ms offset interval, 𝑙*n* corresponds to a natural log transform, and *β*_1_, *β*_2_ are the slope coefficients for spike rate and ISI predictors respectively. Priors were set as broad, uninformative normal priors set on regression parameters. Regression parameters were considered significant if their 95% HDI and credible intervals, summarizing the range in which there is a 95% probability that the true estimate of the regression coefficient lies in the interval given evidence from observed data, does not contain 0. Results are presented as maximum *a priori* estimate of the regression parameters (95% credible interval).

Increases in spike-field coherence as a function of applied INS energy was assessed through Bayesian linear regressions of the form:

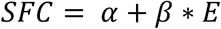

where *α* is the intercept coefficient and *β* is the slope coefficient corresponding to applied natural log INS energy per pulse (𝐸). Observations of SFCs showed that some bands displayed jump discontinuities in regression parameters (Fig. S6), necessitating piecewise linear regression models to capture off vs on states. Piecewise linear regressions were implemented as Bayesian basis spline regression models^95^ of the form:

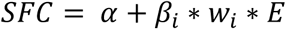

where *α* is the intercept term and *β*_𝑖_ is a spline defined as:

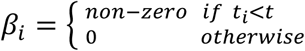

where 𝑡 is the point of concatenation of the piecewise linear regions known as a knot. The 𝑤 term is a learned parameter with the regression slope of each piecewise region described by

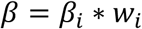

Knot positions were determined by calculating the derivative of the mean SFC as a function of INS energy per pulse and estimating the energy point of a step discontinuity. Knots were confirmed after Bayesian estimation of regression parameters by plotting the domain of each spline to confirm that each spline term only impacts the defined domain above and below the knot. If domain influence significantly overlapped, knots were redefined and models recomputed. Regression parameters were considered significant if their 95% HDI and credible intervals, summarizing the range in which there is a 95% probability that the true estimate of the regression coefficient lies in the interval given evidence from observed data, does not contain 0. Results are presented as maximum *a priori* estimate (95% credible interval) of the regression parameters.

### 2.9 Code, Data, and Materials Availability

Data from this study has been deposited into the following open science framework repository: 10.17605/OSF.IO/JRN65. All source code used in this study is hosted at the following GitHub repository: https://github.com/bscoventry/Focal-Infrared-Neural-Stimulation-Propagates-Dynamical-Transformations-in-A1. A free-standing version of Spyke for LFP analyses can be found in the following GitHub repository: https://github.com/bscoventry/SPyke. Due to online data repository storage limits, raw spike and LFP traces are available upon request.

## 3 Results

### 3.1 Variation of INS Stimulus Parameters Drives Graded LFP Recruitment

To first quantify thalamic recruitment as a function of INS laser parameters, INS dose-response characteristics were modeled using Bayesian hierarchical linear regression (BHLR). BHLR modeled linear changes in LFP RMS voltage between N1-P2 peaks response as a function of INS laser energy-per-pulse, laser ISI, and laser energy-ISI interactions. Regression parameters were summarized by their maximum *a priori* (MAP) estimate (i.e. most probable value) with independent variables considered significant contributors to response if the highest-density interval (HDI) of the parameter distribution corresponding to the 95% most probable parameter values did not overlap 0, following Bayesian inference convention. The 95% HDI credible intervals (CI) are reported to assess uncertainty in parameter estimates. The hierarchical structure of the linear regression model allows for control of both within and between subject variances due to slight differences in electrode placements between animals and active recording channels. All statistical models, sensitivity analyses, and posterior predictive checks are given in the supplementary material. Example subthreshold and suprathreshold LFPs are shown in Fig. 2A. BHLR model responses show that modulation of cortical activity is primarily driven by laser energy per pulse (*β*_1_ = 0.087, 𝐶𝐼: 0.059,0.12) and laser ISI(*β*_2_ = 0.065, 𝐶𝐼: 0.018,0.11). There was no significant interaction between laser energy and ISI parameters(*β*_3_ = −0.0038, 𝐶𝐼: −0.0061,0.0038). Model error was significantly greater than 0, but low in magnitude(∈= 0.51, 𝐶𝐼: 0.47,0.57). The relatively high spread of the energy-per-pulse slope parameter CI suggests a heterogeneity in feedforward thalamocortical activation with increases in laser energy. While increases in LFP magnitude as a function of energy are in line with our previous studies showing that auditory thalamocortical single-unit responses were driven primarily by INS laser energy-per-pulse^3^, the dependence of ISI was not seen in single-unit data.

**Fig. 2.**
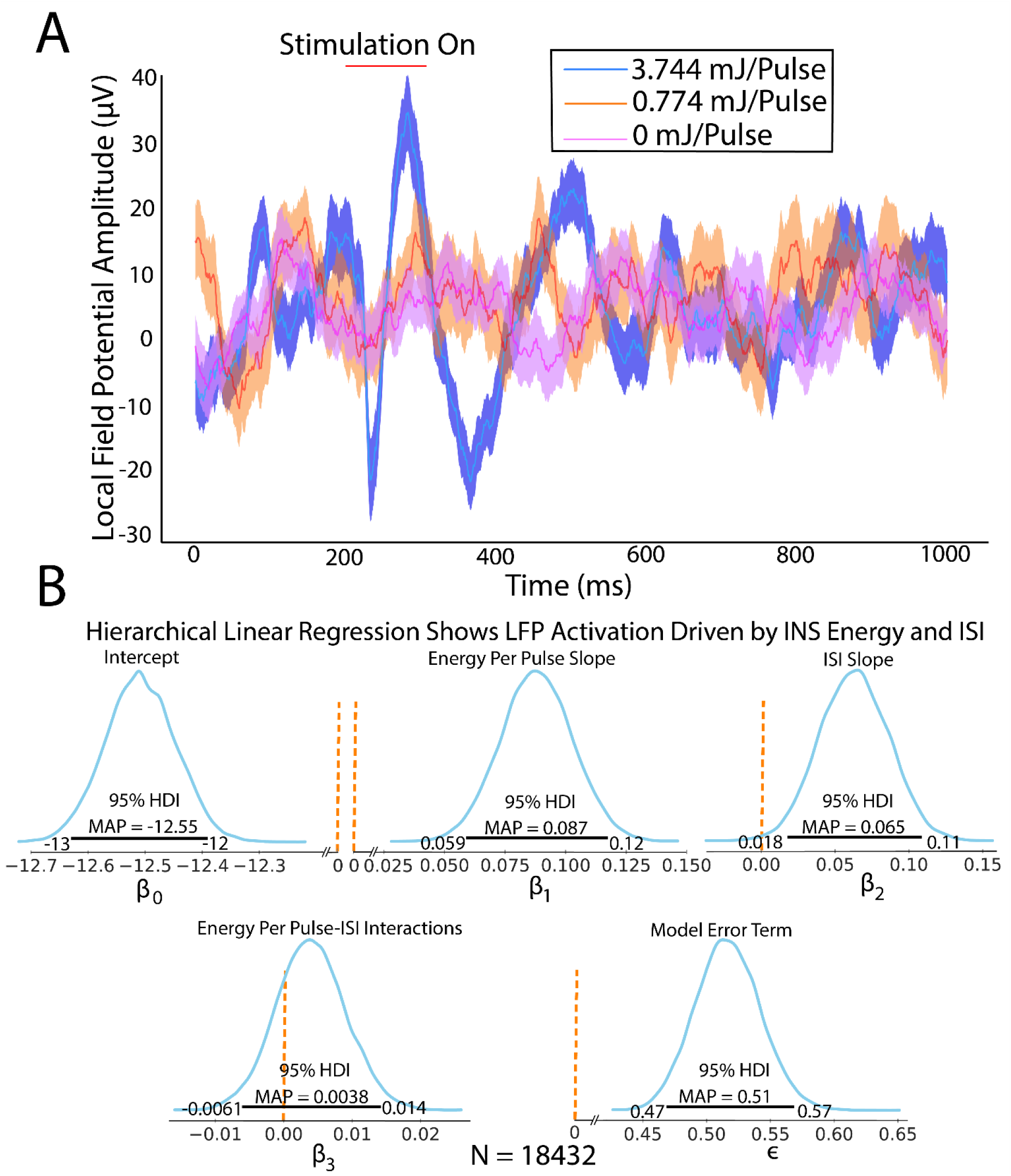
LFP amplitude is driven by increases in INS laser energy-per-pulse and interpulse stimulus intervals. A.) Example LFPs demonstrate increased LFP amplitudes with increased INS laser energy. B.) Bayesian hierarchical linear regression quantifies modulation of LFP N1-P2 RMS voltages. Posterior predictive checks reveal log-log transformed models produced models that best fit observed data. Significance was assessed following Bayesian convention with maximum a priori (MAP) reported. Regression slope parameters were considered significant if posterior distribution 95% highest density interval (HDI) did not contain zero. Credible interval (CI) bounds are given on either side of the HDI boundary. The 0-significance boundary is demarcated by a dotted orange line. Linear regression posterior parameter distributions show that increases in INS laser pulse energy and ISI drive significantly larger N1-P2 amplitudes. There was no evidence of laser energy-ISI interaction. Model errors were significantly above 0 but remained small.

### 3.2 Frequency Band Decomposition

We next quantified the role of thalamic recruitment in modulation of *α*, *β*, *θ*, low *γ*, and high *γ* bands. Time-frequency decomposition of LFPs into constituent frequency bands was performed using continuous wavelet transforms (CWT). A representative example of the LFP spectrogram is shown in Fig. 3A in response to 5 pulses of 3.1 mJ per pulse INS (red line). Pre-stimulus activity was low in all measured frequency bands. INS stimulation elicited a short latency LFP frequency response that showed increased power for frequencies up to about 30 Hz that lasted for hundreds of milliseconds after stimulus offset and approximately 300 ms afterwards. Statistics for the evoked LFP frequency responses are summarized in Table 1. Total band power during onset periods (stimulus window + 100ms) and offset (after onset to end of trial) were calculated. We found significant increases in all frequency bands during stimulation (Table I, Energy slope) as a function of increased energy per pulse. However, band increases were dominated by *β*- band(*β*_*enermmy*_ = 0.17, 𝐶𝐼: 0.089,0.25), low γ-band (*β*_*enermmy*_ = 0.34, 𝐶𝐼: 0.21,0.47, and high γ-band (*β*_*enermmy*_ = 0.49, 𝐶𝐼: 0.32,0.67) with data and regression estimates shown in Fig 3B. There was an observed marginal increase *θ* and *α* LFP band modulation as a function of decreased ISI (*β*_𝐼𝑆𝐼𝜃_ = −0.016, 𝐶𝐼: − 0.031, −0.006, *β*_𝐼𝑆𝐼*α*_ = −0.075, 𝐶𝐼: − 0.1, −0.054). This is indicative of neural integration of multiple pulses into singular impulse events, potentially driving lower frequency onset band responses at lower thresholds. We next examined band activity in the offset period defined as a post stimulation period of 100 ms after stimulus offset to the end of the trial, which ends at 𝑡 = 1 second. Unlike during stimulation, we found significant decreases in *α*, *β*, *θ*, and low *γ* band power from baseline as a function of applied energy (Table I). The high *γ* band, however, did not show any significant INS energy-induced decrease in power. The offset decrease was also most prominent in *β* and low *γ* bands (Fig 3C). This data is consistent with a post-stimulation synaptic inhibition which was also observed in postsynaptic firing in our previous studies^3^. We next assessed the linear covariance between onset and offset *β* − low *γ*, *β* − high *γ*, and low *γ* − high *γ* band powers using Pearson correlation coefficient analysis from responses binned into 500*μ*J bins. Individual frequency bands represent multiscale cortical processing incorporating feedforward excitatory spiking activity, local inhibitory interneuron signaling, and feedback projection activity^96–99^. Pairwise correlations showed increases in positively correlated activity with increased energy per pulse, with *β* − low *γ* showing the strongest positive correlations (Fig 3D). However, the presence of outliers with strong negative correlations suggests that there is heterogeneity in INS driven LFP responses. LFP band correlations in the offset region did not show any pairwise changes in correlated activity (Fig 3E) suggesting covarying band activity is only driven in the INS stimulation window.

**Fig. 3.**
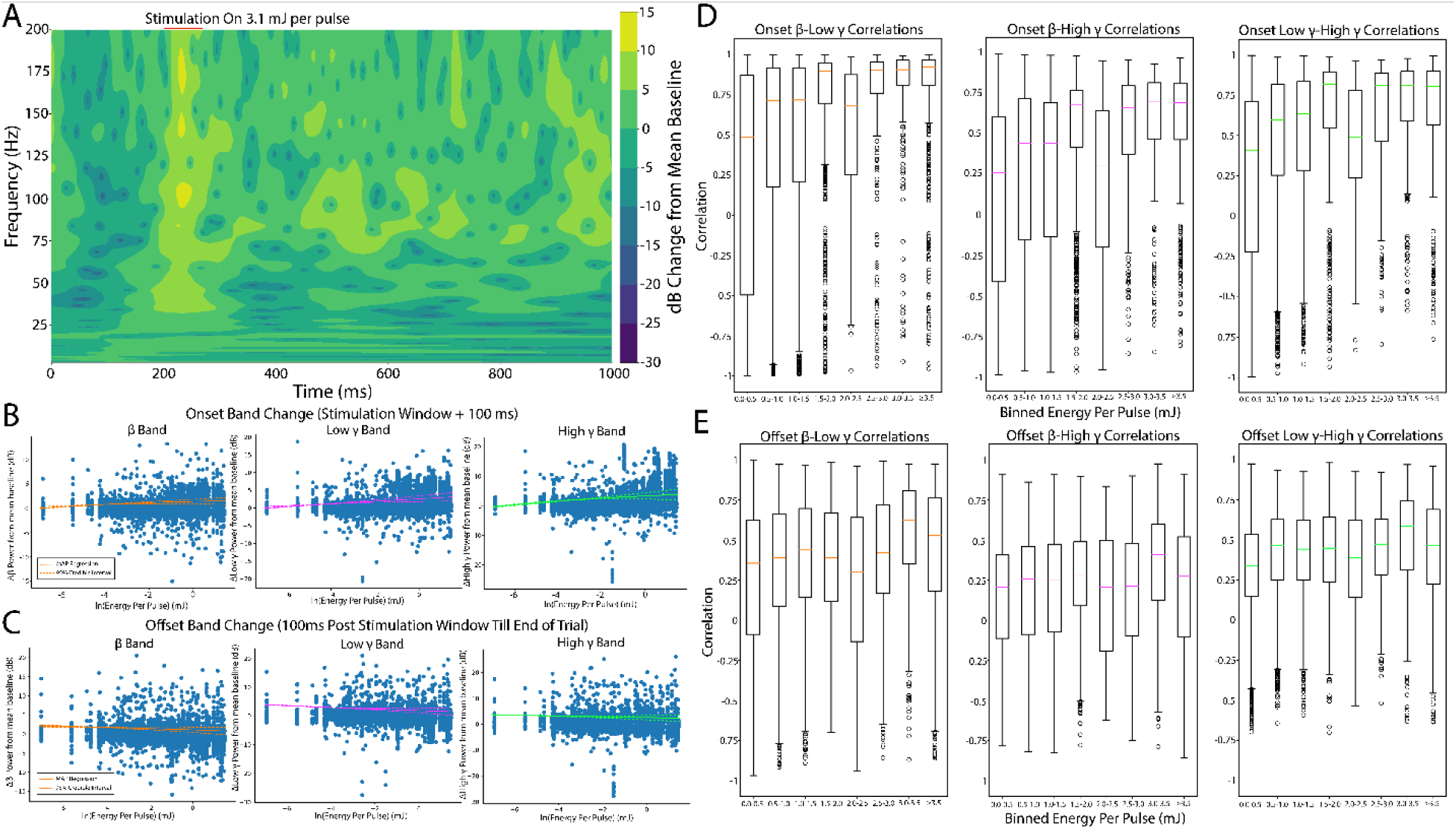
Thalamic INS stimulation predominantly drives activation of *β* and *γ* LFP bands with increases in applied energy. A. Example time-frequency band decomposition calculated using the fast continuous wavelet transform. Analysis was performed by comparing dB changes in evoked LFP band power responses from baseline band power during stimulus onset (time of stimulus on to 100 ms after conclusion of INS stimulus train) and stimulus offset (end of stimulus onset window to end of trial). B. INS activation primarily drives increases in *β*, low *γ*, and high *γ* power bands during onset windows. Responses were graded, showing further power increases with increases in INS energy per pulse. Regression lines are shown as the maximum *a priori* (MAP, solid orange line) with 95% credible intervals (dashed orange line) describing uncertainty in the estimation of the MAP regression. C. Offset responses suggest mild decreases in *β* and low *γ* powers from baseline as a function of applied energy, suggesting a mild post-stimulus inhibition after INS. This was not found in the high *γ* band (MAP slope HDI overlaps 0, Table I). Regression results from all LFP bands for onset and offset windows are summarized in Table I. D. *β*, low *γ*, and high *γ* correlations show increases in between band correlations from baseline and subthreshold activation with correlations generally increasing with increased INS energy per pulse. E. Pairwise offset *β* and low *γ*,and high *γ* correlated band power did not significantly change as a function of applied INS.

**Table I:** Summary results for Bayesian hierarchical linear regressions from models of frequency band decomposition. All results are given as parameter maximum *a priori* estimates (95% credible interval).

|  | Intercept | Energy | ISI | Interaction | Error |
| --- | --- | --- | --- | --- | --- |
| | $\beta_0$ | $\beta_{energy}$ | $\beta_{ISI}$ | $\beta_{E+ISI}$ | $\epsilon$ |
| *Onset $\alpha$* | 0.17<br>(0.13,0.22) | 0.056<br>(0.043,0.067) | -0.075 (-0.1,-<br>0.054) | -0.021 (-0.026,-<br>0.016) | 0.45<br>(0.41,0.51) |
| *Onset $\beta$* | 0.58<br>(0.37,0.79) | 0.17<br>(0.089,0.25) | -0.018 (-<br>0.079,0.015) | -0.017 (-0.025,-<br>0.0066) | 0.89<br>(0.8,1.0) |
| *Onset $\theta$* | 0.037<br>(0.023,0.052) | 0.0093<br>(0.0034,0.015) | -0.016 (-<br>0.031,-0.006) | -0.005 (-0.0087,-<br>0.0027) | 0.24<br>(0.21,0.26) |
| *Onset $\gamma_{low}$* | 1.1 (0.71,1.4) | 0.34<br>(0.21,0.47) | 0.019 (-<br>0.064,0.09) | -0.0021 (-<br>0.018,0.015) | 1.5<br>(1.3,1.6) |
| *Onset $\gamma_{high}$* | 2 (1.5, 2.5) | 0.49<br>(0.32,0.67) | -0.041 (-<br>0.2,0.077) | -0.037 (-<br>0.09,0.0099) | 1.5<br>(1.4,1.7) |
| *Offset $\alpha$* | -0.17<br>(-0.24,-0.096) | -0.074<br>(-0.098,-0.05) | -0.011<br>(-<br>0.034,0.011) | -0.0066<br>(-0.015,0.0029) | 0.84<br>(0.76,0.93) |
| *Offset $\beta$* | -0.23<br>(-0.51,0.036) | -0.14<br>(-0.25,-0.033) | -0.03<br>(-0.0092,0.034) | -0.0074<br>(-0.02,0.01) | 1.4<br>(1.2,1.5) |
| *Offset $\theta$* | -0.049<br>(-0.082,-0.015) | -0.024<br>(-0.038,-0.011) | -0.0062<br>(-0.018,0.0058) | -0.0059<br>(-0.011,-0.00084) | 0.54<br>(0.49,0.6) |
| *Offset $\gamma_{low}$* | -0.078<br>(-0.46,0.3) | -0.24<br>(-0.39,-0.096) | -0.037<br>(-0.1,0.033) | 0.0034<br>(-0.024,0.046) | 2.1<br>(1.9,2.3) |
| *Offset $\gamma_{high}$* | 0.66<br>(0.35,0.97) | -0.1<br>(-0.22,0.0041) | -0.059<br>(-0.14,0.02) | 0.0027<br>(-0.018,0.028) | 2.1<br>(1.8,2.3) |

To evaluate the impact of ISI on LFP entrainment during infrared neural stimulation (INS), temporal modulation transfer functions (tMTFs) were computed. These were defined as the deviation from baseline in ISI frequency power, serving as a metric for the LFP’s synchronization to INS pulses. Results are presented as tMTFs across ISI frequencies and INS energy per pulse bin (Figure 4), with the proportion of LFPs exhibiting significant changes (≥3 dB magnitude from baseline) summarized in Table II. Our findings indicate that LFPs tend to entrain at lower frequencies (longer ISIs) under moderate stimulation conditions (∼2 mJ per pulse). The preferential response to low frequency ISIs at lower energy levels may be explained by shorter ISI being integrated by neurons as singular, longer width pulses. However, increases in INS energy begin to entrain higher frequency ISIs, potentially due to rapid thermal energy gradients which are known to facilitate rapid, low latency responses^29^. However, these trends are not universally represented across all energy bins, with 2-2.5 mJ per pulse bins showing significant responses for 20,40,100 and 200Hz ISIs respectively. Despite this variability, the overall trend of increases in ISI power bands (Table II) suggests that LFP phases generally entrain to INS pulse frequency up to 40Hz and with weaker but significant entrainment even to 100 and 200 Hz stimulation rates. This finding is in alignment with studies showing cortical neuron entrainment to click trains peaking at 50-75Hz repetition rates and weaker entrainment at higher frequencies^100,101^. While more responses showed increases in ISI-band power, some responses demonstrate significant decreases in ISI-band power, indicative of inhibitory entrainment to INS stimulation. This mix of excitatory and inhibitory responses is suggestive of the extensive network modulatory capability of INS. Variability in maximal entrainment frequencies may be further described by exact receptive field placement of recording arrays in A1 which show differential tMTF responses to click stimulation^102^ or the use of dexmedetomidine sedatives which may alter temporal processing^103,104^.

**Fig. 4:**
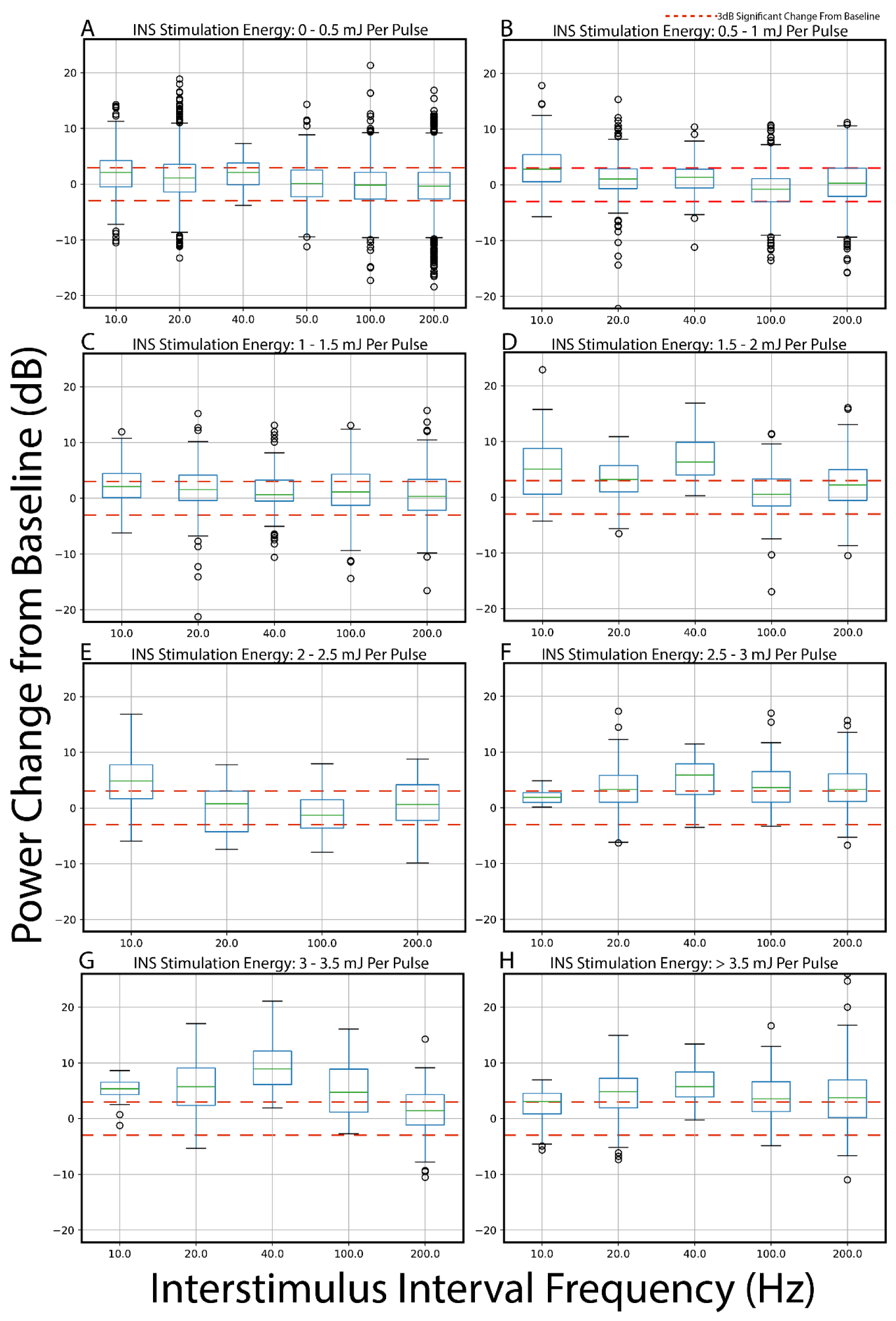
Analysis of INS interpulse stimulation interval tMTFs. Power contained in LFP responses at ISI frequencies was calculated through Fourier analysis with tMTFs calculated as the change of ISI frequency power in stimulation response window from baseline. tMTFs were calculated for binned INS energy per pulse values of A. 0-0.5 mJ, B. 0.5-1mJ, C. 1-1.5 mJ, D. 1.5-2 mJ, E. 2-2.5 mJ, F. 2.5-3 mJ, G 3-3.5 mJ, and H. > 3.5 mJ. Significantly modulated responses are given in Table II.

**Table II:** Percentage of tMTF band power responses above and below 3dB significance threshold.

|  | 10 Hz | 20 Hz | 40 Hz | 100 Hz | 200 Hz |
| --- | --- | --- | --- | --- | --- |
|  | Increased Power |  |  |  |  |
| *0-0.5 mJ/P* | 39.36 | 29.45 | 40.63 | 17.08 | 19.036 |
| *0.5-1 mJ/P* | 48.61 | 22.99 | 23.96 | 13.97 | 25.00 |
| *1-1.5 mJ/P* | 40.63 | 34.38 | 27.08 | 31.77 | 29.17 |
| *1.5-2 mJ/P* | 57.5 | 54.69 | 93.75 | 27.91 | 41.25 |
| *2-2.5 mJ/P* | 66.67 | 25. | N/A | 21.88 | 31.25 |
| *2.5-3 mJ/P* | 18.75 | 55.56 | 71.88 | 54.17 | 51.04 |
| *3-3.5 mJ/P* | 81.25 | 71.88 | 96.88 | 62.5 | 35.42 |
| *>3.5 mJ/P* | 53.125 | 67.19 | 81.25 | 56.25 | 56.77 |
|  | Decreased Power |  |  |  |  |
| *0-0.5 mJ/P* | 9.31 | 14.06 | 3.13 | 22.08 | 21.85 |
| *0.5-1 mJ/P* | 9.72 | 8.71 | 7.29 | 25.37 | 17.63 |
| *1-1.5 mJ/P* | 3.13 | 5.97 | 11.46 | 15.64 | 17.19 |
| *1.5-2 mJ/P* | 7.5 | 4.69 | 0 | 14.84 | 9.58 |
| *2-2.5 mJ/P* | 6.25 | 31.25 | N/A | 37.5 | 17.19 |
| *2.5-3 mJ/P* | 0 | 10.42 | 6.25 | 2.08 | 4.17 |
| *3-3.5 mJ/P* | 0 | 2.34 | 0 | 0 | 13.54 |
| *>3.5 mJ/P* | 12.5 | 3.13 | 0 | 1.04 | 10.42 |

### 3.3 Nonlinear Analysis Reveal Distinct Information Transformation from INS

Previous studies in thalamocortical connectivity have shown that thalamocortical encoding and function is supported by varying states of high and low information transmission capacity which exhibit chaotic or near chaotic dynamics^83^, a state which is called criticality. Criticality may be a feature inherent to normally functioning neural circuits allowing for rapid adaptation to new homeostatic set points from salient stimuli ^105,106^ or as amplification mechanism of synaptic events^107^. To assess the changes in network criticality as a function of INS, the 0-1 test for chaos was used to assess the degree of criticality in a response as a function of the mutual information of stimulus to response. The 0-1 test produces a chaos statistic K describing a measure of “boundedness” of the system, scaled between (0,1), with 0 corresponding to bounded periodic dynamics and 1 corresponding to unbounded, chaotic dynamics. Description and applications of the 0-1 test on canonical nonlinear and chaotic systems can be found in the supplementary methods section (Supp. Fig 1). As the 0-1 test for chaos struggles to delineate true deterministic chaos from stochastic responses, a secondary permutation entropy test for stochasticity^108^ was performed on responses, with only strongly non-stochastic responses included in analysis. After testing for stochasticity, 0.92% of responses exhibited dynamics suitable for criticality measurements. Initial inspection of degree of criticality vs mutual information suggested an inflection point where responses diverged from highly chaotic responses to mixed high and low chaotic responses (Figure 5). Responses were then clustered using K-means clustering into 2 groups, with total number of classes chosen by the Silhouette method^109^. Chaos vs mutual information clustering suggests a fundamental regime change as evidenced by a bifurcation at informative stimulation > 0.4 bits that produces responses which tend toward more deterministic firing as evidenced by lower K-statistic values (Fig 5), suggesting that sub and near-threshold basal firing can exhibit chaotic dynamics. The nature of chaotic dynamics facilitate a high sensitivity to perturbations of the network, which has been demonstrated in computational studies of cortical neurons^110^. Thus, the observed auditory neurons showing chaotic responses may allow for rapid adaptation to temporally complex stimulation^111^ which may not be necessary when highly informative stimulation activates well-defined thalamocortical networks. This transition to purely deterministic dynamics is not complete, however, as there exists a subset of responses which maintain highly chaotic firing patterns in tandem with high mutual information. This state represents responses which are separable by applied INS energy which still exhibit drastically varying response profiles, and thus suggests field-potential responses which are continually at the edge of criticality. This potentially is a result of groups of neurons which naturally exhibit chaotic dynamics as part of normal cellular processing by facilitating the encoding of rapidly changing sensory^112^ or synaptic^107,113^ inputs.

**Fig. 5:**
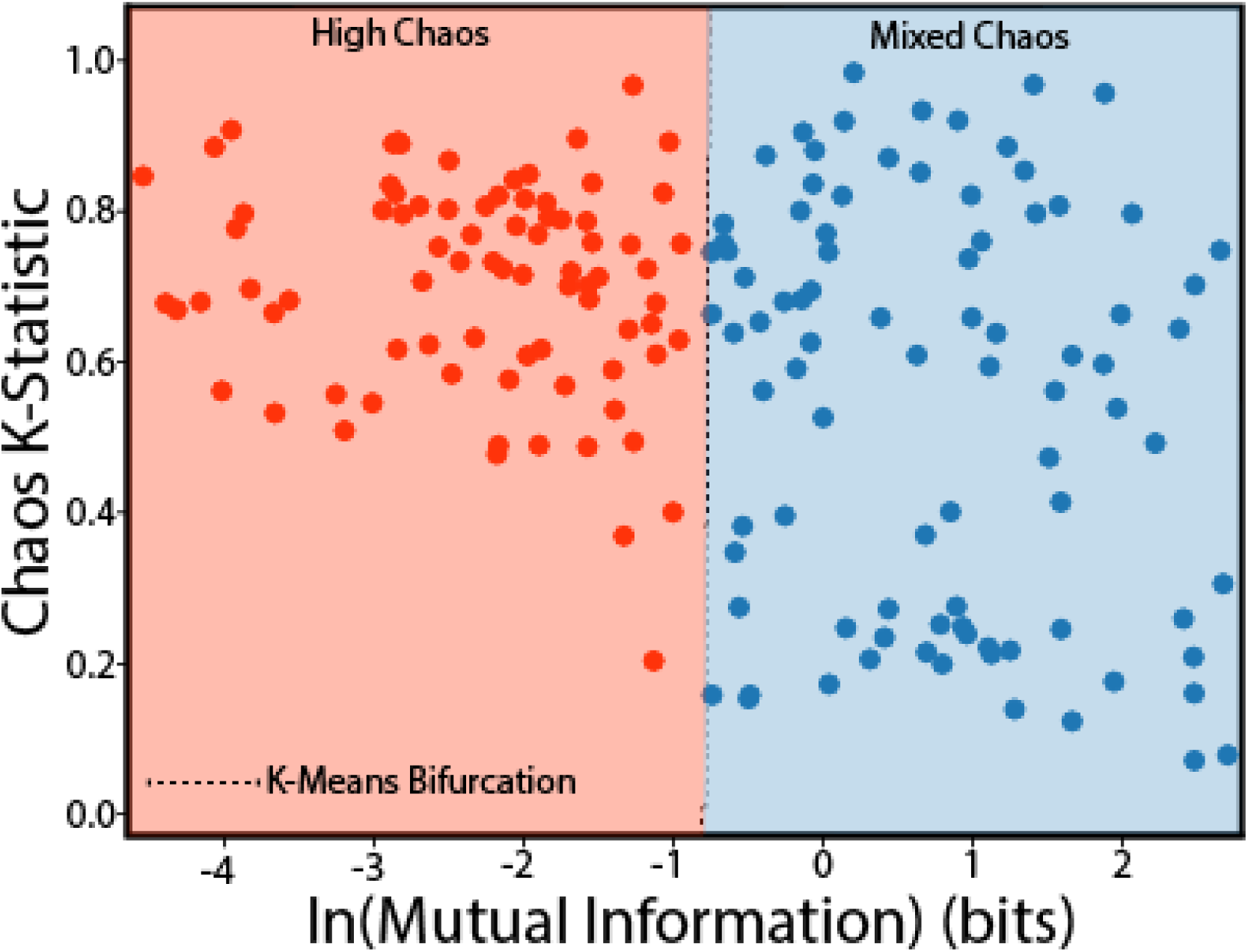
Observation of chaotic dynamics in LFP responses is correlated to stimulation to LFP information transfer. 0-1 Chaos tests were used to assess the criticality of recorded LFPs and plotted vs mutual information of stimulus to response. Silhouette methods were used to determine the optimal number of K-means clusters (K=2). Results show a bifurcation point at stimulus-response mutual information > 0.4 bits suggest a fundamental firing state changes for neurons displaying critical, chaotic dynamics at baseline to mixed chaos responses. Mixed chaos responses are suggestive of temporally precise adaptation to changing sensory stimulation.

### 3.4 LFP Amplitudes are Correlated with Evoked Spike Rates

We next investigated the correlation between driven single unit responses (stimulation spike rate≥ 1.75 basal firing rate) and LFP N1-P2 RMS amplitudes. Bayesian linear regressions (Fig 6 A,B, Bayesian P-value Fig. S5) suggest that LFP N1-P2 RMS amplitudes increase with increased natural log-transformed spike rate (*β*_𝑠𝑝𝑖𝑘*er*𝑎𝑡*e*_ = 1.8, 𝐶𝐼: 1.7,2) and decreased RMS amplitudes with increased natural log-transformed ISI (*β*_𝐼𝑆𝐼_ = −0.047, 𝐶𝐼: −0.059, −0.037), suggesting sublinear LFP summation for short-interval pulses. Comparing responses from fixed ISIs (Fig 6 C-E) suggested that short ISI pulse trains integrate to produce higher amplitude N1-P2 LFP and spike rate responses while longer ISI trains produced lower LFP and spike rate responses indicated of discrete pulse events driving moderate responses. Similar spike-LFP synergies have been observed in sensory regions across cortex in response to sensory stimulation^114–116^ and is suggestive that INS can elicit sensory relevant responses in cortex.

**Fig 6:**
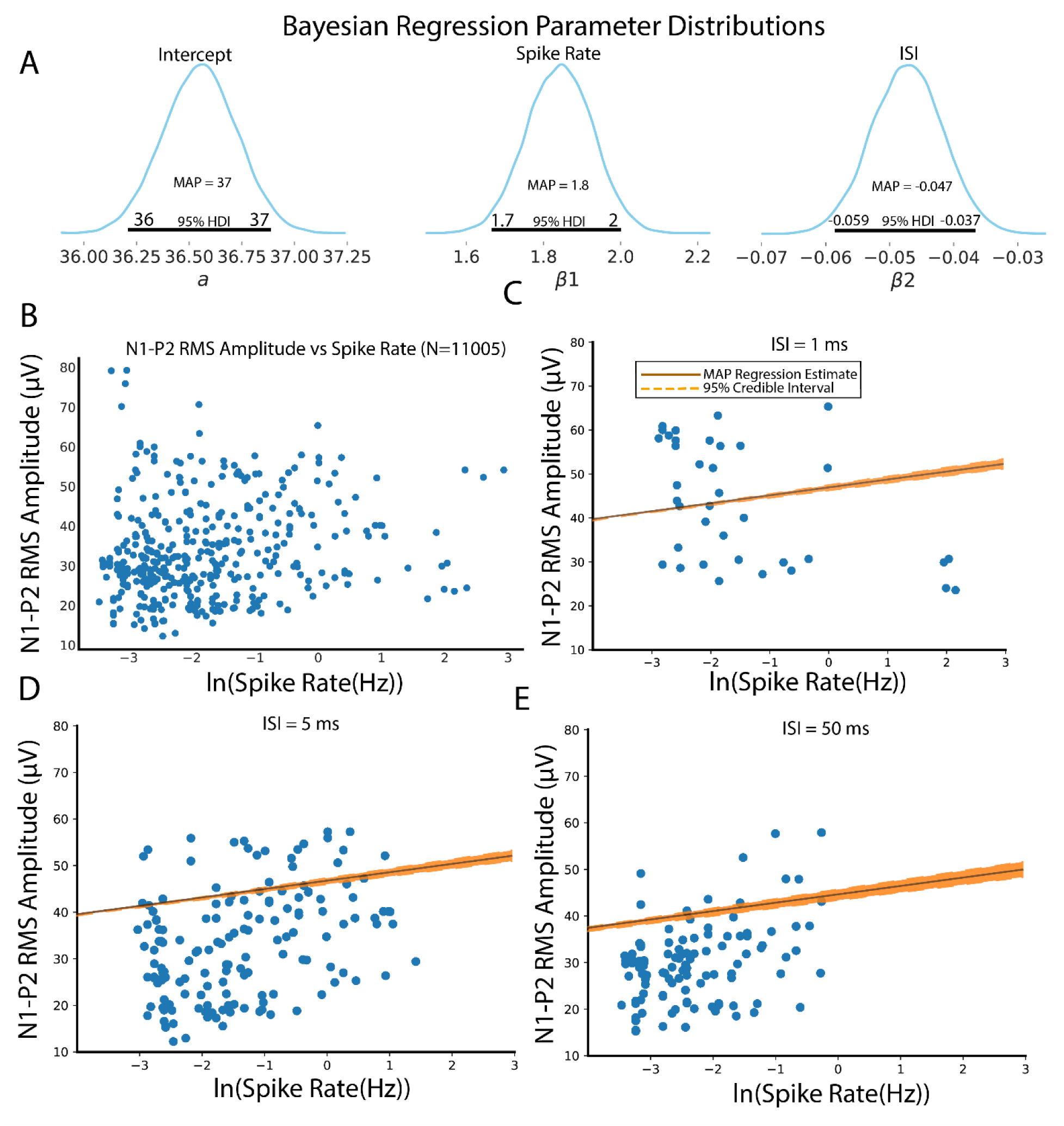
Spike rates vs LFP N1-P2 magnitude correlations. A.) Regression parameter distributions suggest a significant intercept term greater than zero with LFP N1-P2 RMS amplitudes showing increases with log spike firing rates and decreases with log ISI. B.) Scatter plot of spike rate and N1-P2 LFP RMS amplitudes across all ISI values. C-E.) Regressions for spike rate, N1-P2 amplitude correlations for 1ms ISI (C), 5 ms ISI (D), and 50 ms ISI (E). Regressions are reported as MAP parameter estimates with shaded regions representing the 95% credible interval.

### 3.5 Dual Regime Synaptic Transformations Are Driven by INS

Firing of thalamocortical neurons, whether driven by sensory, motor, cognitive, or INS stimulation, induces a rapid reorganization of local cortical activity due to spiking and synaptic activities^45,117,118^. We utilized spike-field coherence (SFC), a measure of system causal input-output power transfer, as a function of applied INS energy to assess frequency-band specific network modulation. Dependence of spike-LFP coherence on INS laser parameters was then assessed using Bayesian multilinear regressions, with all results presented in table III. SFC regressions are shown in Fig 7A. Regressions revealed a slow increase in high *γ* SFC (*β_himmh γ_* = 0.015, 𝐶𝐼: 0.015,0.016) in response to log increase in INS energy. Increases in *γ*-band SFC are linked to faster afferent spiking activities^98,119^ as well as parvalbumin(PV)-expressing interneuron activities^120^, suggesting that increased INS energy drives increased LFP to spiking activity with putative interneuron recurrent inhibition, similar to what has been seen in electrically mediated neural recruitment^121^. This is also consistent with our previous results showing log-linear increases in spiking activity with increased INS energy^3^. Studies utilizing electrical stimulation of auditory thalamocortical circuits display similar interlayer side-band inhibition^122^ which can putatively be attributed to interneuron inhibition. Observation of low *γ* regressions suggested the presence of a nonlinear discontinuity in spike-field coupling (supplementary figure S6), necessitating a piecewise linear regression model. Low *γ* SFC dynamics were captured using a basis-spline regression model performing piecewise linear estimates around the nonlinear jump discontinuity, called a knot. The low *γ* knot was determined by estimating the step discontinuity in the derivative of the mean SFC as a function of ln(energy), with knot placement confirmed by observation of the spline domain to determine relative influence of each spline in the total regression model. Piecewise regression coefficients are shown in Table III. Low *γ* knot was determined to occur at 0.15 mJ per pulse, below which spike-field coupling showed no significant increase as a function of applied energy per pulse (*β*_𝑙𝑜*ee* *γ*_ = 0.048, CI: 0,0.07). However, above the knot value, low *γ* spike-LFP coupling significantly increases as a function of INS energy (*β*_𝑙𝑜*ee* *γ*_ = 0.156, CI: 0.11,0.262). The presence of a discontinuity suggests an early local activation followed by a rapid increase in the number of cortical afferents recruited. SFC regressions in the *β* band also suggested the presence of a step discontinuity (supplementary figure S4). The knot point in *β* SFCs was determined to be at 0.125 mJ per pulse. Piecewise basis spline regressions showed no significant increase in *β* band SFC (*β*_*β*_ = 0.034, CI: 0,0.05) as a function of INS energy below the knot and a significant increase in *β* band SFC (*β*_*β*_ = 0.143, CI: 0.0979, 0.24) as a function of INS energy above the knot. Synchronous *β* coherence is associated with multi-synapse thalamocortical feedback activity^123^ and a broad range of interneuron subtypes^97^. Finally, linear increases in *α*-SFCs were observed (*β*_*α*_ = 0.024, CI: 0.023,0.024), even at low stimulus energies *α*-SFCs have been associated with stimulus adaptation^124^ also from thalamocortical feedback circuits and cortical inhibitory processing^96^. It should be noted that a small, but significant increase in *θ*-SFC was also observed(*β*_𝜃_ = 0.0037, CI: 0.0034,0.004). However, given the small regression MAP estimate, this increase is likely due to low frequency LFP drift coupled with co-activation by INS.

**Fig. 7:**
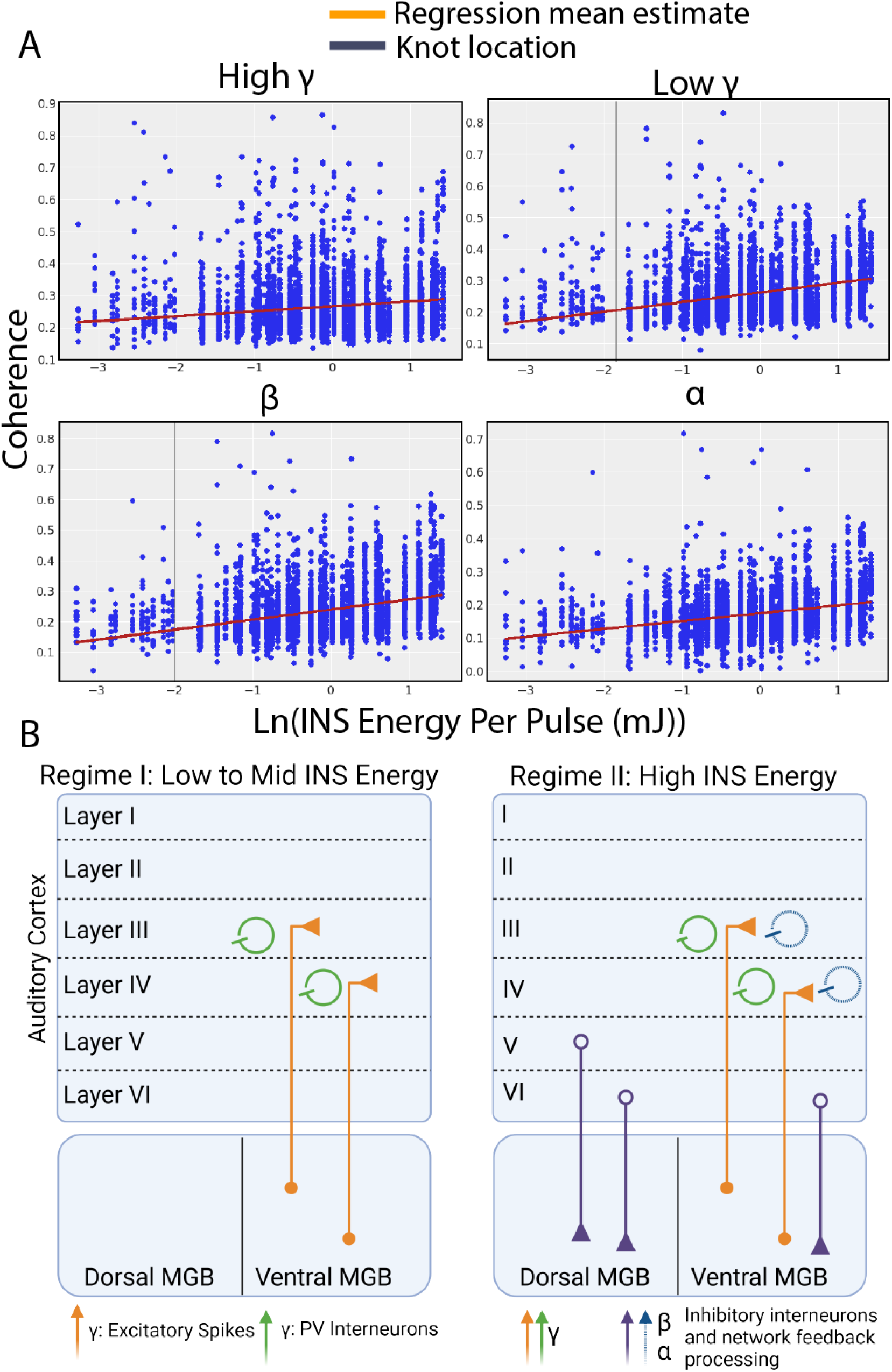
Spike-field coherence (SFC) regressions create a model of network entrainment from INS stimulation. A. *α*, *β*, and *γ* SFC regressions as a function of INS energy per pulse. All regression fits represent the mean estimate of regression parameters, with MAP parameter values and 95% HDIs given in table III and IV. SFCs in *α* and high-*γ* bands show linear behavior as a function of natural log increases in INS energy per pulse. SFCs in the *β* and low-*γ* band show log-linear behavior in higher energies which deviates at lower energies, suggestive of a nonlinear jump discontinuity. To account for this discontinuity, piecewise linear spline regressions with 1 joint knot were performed. *β* SFCs showed no significant increase below INS energies of 0.125 mJ with linear increases after 0.125 mJ. Similarly, low *γ* activity showed no significant increases in coherence below INS energies of 0.15 mJ per pulse with log-linear increases in response to increases in INS energy above 0.15 mJ. Vertical grey lines denote the point of knot discontinuity. B. Graphical model of INS network recruitment of the auditory thalamocortical circuit. INS recruitment of the auditory thalamocortical circuit is characterized by two distinct regimes: a low energy regime characterized by thalamic recruitment of Layer III/IV excitatory projections and PV-interneurons and a high energy regime characterized by further excitatory recruitment with recruitment of Layer V/VI feedback projections and interneurons of all types.

**Table III:** Spike-field coherence regression model parameters for *α* and high *γ* bands.

| | Intercept<br>($\beta_0$) | Energy ($\beta_{energy}$) |
| --- | --- | --- |
| $\theta$ | 0.185 (0.1843, 0.1847) | 0.0037 (0.0034, 0.004) |
| $\alpha$ | 0.17 (0.1738,0.1744) | 0.024 (0.023,0.024) |
| $\gamma_{high}$ | 0.2656 (0.2652, 0.2662) | 0.015(0.015,0.016) |

**Table IV:** Spike-field coherence regression model parameters for *β* and low *γ* bands.

| | Intercept<br>$(\beta_0)$ | Energy Slope Low<br>$(\beta_{energy})$ | Energy Slope High<br>$(\beta_{energy})$ |
| --- | --- | --- | --- |
| $\beta$ | 0.034 (0.0247,0.0249) | 0.034 (0,0.05) | 0.143 (0.0979, 0.24) |
| $\gamma_{low}$ | 0.2656 (0.2652, 0.2662) | 0.048 (0,0.07) | 0.156 (0.11,0.262) |

This data thus suggests two separate thalamocortical encoding paradigms: a first state in which low to moderate INS stimulation drives weak local activity in cortex amplified by cortical feedback expressed as low level *α* activity with log-linear increases as a function of stimulus energy. Increased INS stimulation energy then causes a shift to a second state characterized by onset of *β* and *γ* activity, with increases in INS energy leading to log-linear increases in these bands. Given the observed data and extensive knowledge of auditory thalamocortical architectures, we propose a model of thalamocortical INS recruitment, schematized in Fig 7B. INS at near threshold energy levels recruits local excitatory projections^125^ driving local activation subthreshold to observation but facilitated and amplified by *α* band cortico-cortical feedback^126^. Increases in INS energy per pulse will eventually cross a threshold in which substantial spiking activity is recruited^3^, evidenced by increased *γ* band activity and supported through *α* and *β* band cortico-cortical feedback projections with PV interneuron activity concentrated in layer III/IV. We suggest that the second regime is characterized by more widespread activation of interneurons, including PV and somatostatin expressing subtypes^127^. It is also likely that increased stimulation energies will further drive cortical-thalamic and cortical-fugal feedback projections from layer V/VI, but these projections likely only weakly drive observed cortical LFP activity in lower frequency *α* and *β* bands.

## 4 Discussion

While INS is increasingly used in the study of neural circuit function and evaluated for clinical neuromodulation therapies, the stimulus-driven dynamics for thalamocortical INS has, until this point, not been a primary focus of study. Surface or intracortical stimulation has typically been used to drive cortical activity, rather than the natural thalamocortical pathway^11,12,17,128^. One prior thalamocortical INS study other than our lab examined slower thalamocortical connectivity using fMRI in anesthetized macaques^44^. In this study, we evaluated thalamocortical encoding of INS stimulation using measures of local field potentials as a readout for cortical network modulation resulting from thalamic stimulation in a chronic rat preparation.

### 4.1 LFP Magnitude is Driven by INS Stimulation Energy and ISI and is Coupled Primarily to β and γ LFP Frequency Bands

We tested the relationships between stimulus-intensity and frequency band-specific LFP magnitudes, which is a crucial characteristic for basic or translational use of INS. We showed that cortical network dynamics are driven by applied INS energy (Fig. 2) and have significant entrainment to stimulus pulse frequencies even up to 200 Hz (Fig. 4). Notably, our results indicate that cortical temporal encoding of INS stimulation frequencies followed patterns similar to those observed in auditory click train processing. Specifically, the temporal modulation transfer functions (tMTFs) of LFP responses to INS resemble those elicited by auditory click train stimulation^100,129^. This data suggests that INS can drive standard temporal coding in cortical neurons with a mechanism of activation sufficiently fast enough to recapitulate thalamocortical temporal responses.

Decomposition of the LFP into frequency bands also showed that INS activity is primarily coupled to *β* and *γ* band activity. Specifically, INS energy per pulse couples strongly to *β* and high-*γ* frequency bands with weaker coupling to *α* and low-*γ* bands. Furthermore, *α* and *β* bands are significantly increased with decreased ISI, suggesting that INS pulses close in time can integrate to create stronger activation while pulses with longer ISIs create distinct activation events that are adapting to multiple pulses. Similar high-*γ* coupling may also arise from thalamic rebound firing to stimulus offset^130–133^. Delineation of thalamic tonic vs rebound firing could not be assessed in the present study but should be investigated in future patch clamp studies. INS coupling to *β* and *γ* bands has interesting clinical potential, with modulation of these bands used as physiological readouts for closed-loop deep brain stimulation efficacy in Parkinson’s disease^66,134,135^ and obsessive-compulsive disorder^136,137^. Whether or not similar entrainment is consistent between similar thalamocortical and subcortical-cortical circuits related to these diseases requires further study.

### 4.2 Neurons Showing Chaotic Dynamics Exhibit Distinct Firing State Changes from INS Stimulation

Linear methods of assessing neural encoding of artificial and sensory stimulation are powerful; but fundamentally limited to studies of neural activity with strong linear entrainment to a stimulus. While linear methods are often sufficient for the understanding of neural coding, nonlinear approaches can provide a deeper understanding of spontaneous and evoked neural behavior and activity. In this study, we used the 0-1 test for chaos to assess criticality in the auditory thalamocortical circuit. As chaotic dynamics in other thalamocortical circuits were found to correlate with information transfer capabilities^83^ coupled with INS’ ability to produce information- rich responses in auditory cortex^3^, we mapped statistics indicative of chaotic dynamics to thalamocortical information transfer from INS stimulation and found evidence of chaotic activity in the auditory cortex that is most prominent in subthreshold stimulation in a subset of A1 neurons (Fig. 5). While we cannot properly deduce the primary function of chaotic firing in this study, chaos in other cortical circuits has been linked with facilitation of information transmission^138^ and entrainment to temporally complex firing^112^, features of which may be critically important in A1 function. We further show a bifurcation in chaotic firing states from complete chaotic firing regimes to mixed chaotic and non-chaotic responses that were correlated with information transfer.

This suggests that some neurons display chaotic firing during subthreshold or low-level stimulation that converge to more stereotyped, linear responses when stimuli drive informative thalamocortical responses. These responses may act as adaptive feature detectors to facilitate dynamic range adaptation present in subsets of A1 neurons^46,139,140^. However, other responses observed exhibit chaotic dynamics across all stimulus information sets. This subclass of neurons display constant chaotic dynamics may be a fundamental feature of A1 neural circuits to facilitate rapid circuit gain changes and pattern detection^112^ capabilities. It is likely, however, that the number of neurons expressing chaotic dynamics that we observed is undersampled, as passing the permutation entropy test for stochasticity was a stringent requirement for inclusion in chaos analysis. This precludes responses which may have mixed deterministic chaos and stochastic responses, in which random dynamical systems theory^141^ may be used to better understand chaotic dynamics in the presence of stochastic dynamics. Furthermore, the study of stochastic dynamics and chaos is still in its infancy^142^ with others suggesting that the chaos seen in neural circuits represents an entirely new form of dynamical process^143^. We also believe that similar INS studies can be used to better understand neural dynamical processes as INS produces no electrical artifact into evoked responses and is easily used in both *in vivo* and *in vitro* preparations.

### 4.3 Thalamocortical Transformations Underlie The Encoding of INS Stimulation

SFC regression analysis revealed two distinct thalamocortical regimes of dynamic thalamocortical transformations in response to INS. The first regime is characterized by focal excitation of primarily excitatory afferent responses as evidenced by increases *γ* SFC indicative of thalamocortical synapses onto dendrites driving spiking activity as well as PV interneuron activity^98,119,120^. The first regime also has evidence of slight stimulus driven cortico-cortico feedback, as evidenced by slight increases in *α* SFC^124^. The second regime is characterized by full entrainment of the thalamocortical circuit with continued cortico-cortical feedback excitatory activation and PV interneuron activation as evidenced by *γ* SFC along with recruitment. Auditory *β* and high *γ* coupling also reflects stimulus evoked synaptic drive localized to microcircuit activation^98^ potentially from neurons with similar receptive fields^144^. Other evidence in visual cortex also suggests that *β* and high *γ* coupling involves single and multisynaptic coupling^123^. It should be noted that *α* and *β* power bands can also reflect cortico-thalamo-cortical feedback loops as a part of normal function of thalamocortical processing^145,146^. Our current study was not designed to assess corticothalamic feedback, which should be investigated for full understanding of INS stimulation of thalamocortical circuits. Our data provides a model of network thalamocortical activation (Fig 7B) which can help facilitate understanding of the therapeutic potential of INS. INS intensity, pulse characteristics, and pulse timing as variables generates a vast parameter space for control of thalamocortical dynamics, including frequency-band specific modulation that may be relevant for restoring function or managing pathological conditions. This is particularly salient for the clinical use of INS in which a myriad of network recruitment profiles may be desired^147,148^. We also note that the thalamocortical order of recruitment presented by our model is in alignment with naturalistic thalamocortical recruitment in somatosensory^149,150^ and auditory cortices^127,151,152^, in response to naturalistic stimuli, as well as in human thalamocortical circuits in response to propofol-induced brain state modulation^153^, further suggesting INS drives naturalistic neural firing. It is also likely that thalamocortical and activation will also recruit thalamic reticular network neurons^154^ coupled to *α* and *β* oscillations^145,155^. This is not yet included in our model as thalamic reticular network neuronal responses to thalamic INS were not measured.

### 4.4 Towards a Full Coverage Auditory Thalamic Neuroprosthesis

While the cochlear implant (CI) remains one of the most clinically successful neuromodulation tools, there remains a dearth of options for patients who cannot benefit from CI use. One example is that of patients diagnosed with Neurofibromatosis Type 2, characterized by tumor growth on the auditory nerve^156^. While this patient group may receive auditory brainstem implants, the tumor resection surgery is known to damage the cochlear nucleus implantation site^157^, limiting the efficacy of therapeutic stimuli^158^. While inferior colliculus auditory midbrain implants have also been tested^159,160^, the inferior colliculus is a difficult implantation target due to the need to replicate the convergence of disparate inputs from all lower auditory nuclei^161–163^. The MGB, however, is characterized by a small number of parallel sets of excitatory and inhibitory inputs from inferior colliculus to MGB^50,155,164,165^, with little long term potentiation or depression after development^166^, potentially allowing for the generation of robust stimulation waveforms to generate reliable auditory percepts. Furthermore, the MGB is adjacent to areas commonly targeted by deep brain stimulation^167^, so it is more straightforward to access surgically. The present INS encoding data combined with INS’ advantageous spatial specificity, information transfer properties^3,10,14,22,168^ and favorable safety profile^16–19,22^ suggests that an INS-based thalamic implant may serve as a potent auditory restoration tool. An example of such a device is shown in Figure 8 for human cortex^167^. Much progress has been made in fabricating optical devices for chronic stimulation^20,169,170^, with the potential for multicolumnar stimulation^35^ facilitating the creation of stimulation devices similar to clinical gold standard stimulation arrays. One potential implementation (Fig.8 top) would be a coronal plane implantation, which would allow for recruitment of primary (ventral MGB) and non-primary (dorsal and medial MGB). A potentially more advantageous approach would be a sagittal-plane implant (Fig 8 Bottom), allowing for more optrode coverage in the ventral division of MGB and the potential to provide finer resolution of input auditory waveforms. Some coverage of nonlemniscal dorsal MGB can also be leveraged to modulate circuit state and large-scale gain control and modulation of auditory responses^155,165,171,172^, providing a pathway for precise control of lemniscal and nonlemniscal auditory function. However, the amount of MGB coverage for restoration to cochlear-implant level of hearing is not known, requiring fine-tuned behavioral and information processing studies in preclinical animal models. Furthermore, INS neural recruitment, like conventional electrical stimulation, is heavily dependent on the geometry of the stimulated area. As INS activation profiles are dependent on the width of the optical stimulation area and the penetration depth of the chosen wavelength^15,173^, the total volume of tissue recruited in small animal models is likely to be larger than what is observed in human. Therefore, larger animal studies combined with computational modeling of optic field propagation in human MGB is necessary to help establish best electrode design for implanted auditory stimulation optrodes.

**Fig. 8:**
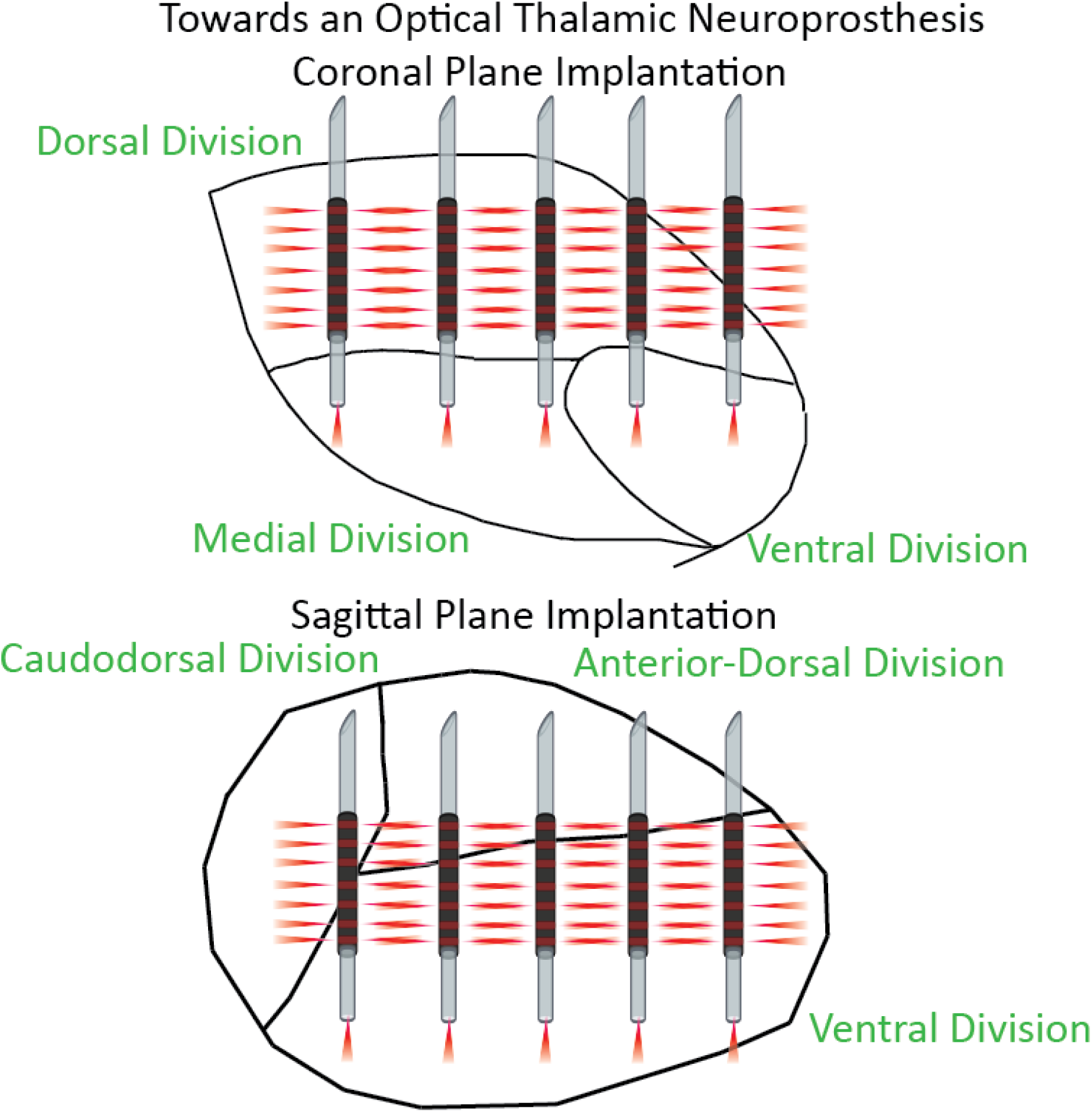
A proposed full coverage INS-based thalamic neuroprostheses. A. Coronal implantation plane provides minimal coverage of ventral MGB with larger coverage of nonlemniscal auditory regions. B. Sagittal implantation routes would provide large INS coverage of lemniscal auditory pathways with adequate coverage of nonlemniscal regions which can either remain unused or serve as a route to account for auditory modulation and control.

### Disclosures

BSC is an unpaid scientific consultant for BECATech Inc for work unrelated to the present study. BSC also holds provisional patents in neural spectral decomposition methods. The other authors declare no conflicts of interest for the data presented in this study.

### Code, Data, and Materials

The data and statistical posterior distributions presented in this article are publicly available in an Open Science Framework repository at 10.17605/OSF.IO/JRN65. All source code for data analysis and statistics is archived in a GitHub/Zenodo repository at https://github.com/bscoventry/Thalamic-Infrared-Neural-Stimulation-Propagates-Linear-And-Chaotic-Coding-Transformations-in-A1. Build instructions and bill of materials for the INSight optical neuromodulation platform can be found in the following Zenodo repository: 10.5281/zenodo.11086673

## Acknowledgments

This study was supported by grants from the National Institutes of Health (NIDCD R01DC011580, PI: E.L.B.) and the Purdue Institute for Integrative Neuroscience collaborative training grant (PI: B.S.C.). BSC is now with the Wisconsin Institute for Translational Neuroengineering, the Department of Neurological Surgery, and the Department of Biomedical Engineering, University of Wisconsin-Madison USA.

**Brandon S Coventry** received his BS in electrical engineering from Saint Louis University, his MS in electrical and computer engineering from Purdue University and a PhD in biomedical engineering from Purdue University. He is currently a postdoctoral research associate at the Wisconsin Institute for Translational Neuroengineering and the Department of Neurological Surgery, and a lecturer in the Department of Biomedical Engineering, University of Wisconsin-Madison. His current research interests include translation of clinical neuromodulation tools, deep brain stimulation, infrared neural stimulation, and thalamocortical circuit function.

**Cuong P Luu** received his BS in bioengineering from UC Berkeley. He is currently a medical student at the University of Wisconsin, School of Medicine and Public Health. His current research interests span deep brain stimulation, neuromodulation tools, and brain computer interfaces in addition to the translation of discoveries in these fields into functional neurosurgery.

**Edward L Bartlett** received his BS in physics from Haverford University and PhD in neuroscience from the University of Wisconsin-Madison. He is currently a Professor of Biological Sciences and the Weldon School of Biomedical Engineering and Associate Dean for Undergraduate Affairs, College of Science, Purdue University and is an affiliate of the Purdue Institute for Integrative Neuroscience. His current research interests include dynamic interactions between auditory thalamus and cortex, alterations in hearing function from noise exposure and aging, and infrared and closed-loop control of auditory circuits.

## Supplemental Material Outline

### 1. Visualization of 0-1 Chaos Testing

The 0-1 test for chaos is a method by which the degree of chaos in deterministic dynamical systems can be determined without the need for phase space reconstruction which is difficult if not impossible for systems without explicit mathematical models. As such, the 0-1 test can act as a surrogate for Lyapunov exponent methods^83^. The 0-1 chaos test operates through mapping dynamical time series to a 2D system:

The 0-1 test for chaos is a method by which the degree of chaos in deterministic dynamical systems can be determined without the need for phase space reconstruction which is difficult if not impossible for systems without explicit mathematical models. As such, the 0-1 test can act as a surrogate for Lyapunov exponent methods^83^. The 0-1 chaos test operates through mapping dynamical time series to a 2D system:

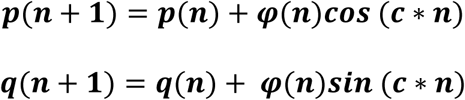

and then computing the mean path displacement as

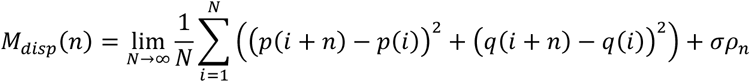

The measure of degree of chaotic dynamics is then summarized in the K-statistic

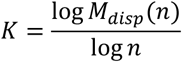

The result is that non-chaotic dynamics display periodic, bounded dynamics in 𝑝(*n*), 𝑞(*n*), bounded growth 𝑀_𝑑𝑖𝑠𝑝_(*n*) and a resulting K statistic approaching 0. Alternatively, chaotic systems display diffusive, Brownian-like motion in 𝑝(*n*), 𝑞(*n*), unbounded linear growth in 𝑀_𝑑𝑖𝑠𝑝_(*n*) and a K-statistic approaching 1. To visualize test dynamics, the logistic map was used:

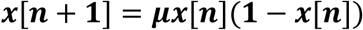

Where μ is known as the growth rate. Plots of the logistic map as a function of *μ* are shown in FigS1(top). Values of *μ* ≤ ∼3.5 show nonchaotic dynamics while values greater than this critical threshold show chaotic, period-doubling behavior. Plotting 𝑝(*n*), 𝑞(*n*) at *μ* = 2.8 displays sinusoidal, bounded behavior (FigS1, bottom left) with a resulting K-statistic of 0.1789, indicative of non-chaotic dynamics. Plotting 𝑝(*n*), 𝑞(*n*) at *μ* = 3.8 displays Brownian motion-like behavior (FigS1, bottom right) with a resulting K-statistic of 0.9982, indicative of chaotic dynamics. Full mathematical descriptions and validations of the 0-1 test for chaos can be found in the following studies^81,174–177^.

### 2. Bayesian Model Descriptions and Sensitivity Analyses

This study follows the guidelines outlined in the best practices in reporting Bayesian analysis^93^, and is composed of:

- Descriptions of necessary software for replication
- Depositing of posterior traces into freely available repositories
- A description of the goals of analysis
- Detailed model descriptions
- Detailed description of prior distributions and reasoning behind choice of priors
- Sensitivity Analysis for statistical models
- Posterior and Markov-chain Monte Carlo diagnostics

#### 2.1 Software requirements

The Bayesian inference performed in this study was computed on intel processors with no additional GPU threading. Replication of Bayesian models is CPU bound which can be performed on most current generation CPU platforms. All models where implemented in the PyMC environment^178^ running in Python 3. All model source code is available on our Github repositories for this article (data and code availability) which can reproduce posterior traces used in this study. Posterior traces can also be made available upon request.

#### 2.2 Goals of Bayesian Analysis and Inference

Bayesian inference is particularly well suited for inference of repeated measures study designs which contain within and between subjects variance through flexible hierarchical inference structures^89,179,180^ allowing for quantification of heterogenous LFP responses from collective activity of groups of biophysically differing cell types^181^. Bayesian inference is particularly well suited for inference of repeated measures study designs which contain within and between subjects variance through flexible hierarchical inference structures^89,179,180^ allowing for quantification of heterogenous LFP responses from collective activity of groups of biophysically differing cell types^181^. As such, Bayesian hierarchical linear regressions were used to quantify dose-response characteristics of INS-LFP entrainment modeling increases in wave N1-P2 peak RMS as a function of INS energy per pulse, INS interpulse stimulation intervals (ISI), and energy-ISI interactions. Models also contained error term parameters to quantify distributions of fitting errors. Changes in LFP *θ*, *α*, *β*, and *γ* band powers were also quantified using Bayesian hierarchical linear regression models with INS energy per pulse, INS interpulse stimulation intervals (ISI), and energy-ISI interaction predictors. Hierarchical models perform ‘partial pooling’ of response data which accounts for individual differences in parameter estimation. This is done by allowing distributions around model that quantify firing rate dependencies from laser parameters while accounting for within-subject differences. Prior selection is discussed in section 2.3. Partial pooling was performed by adding an implicit class definition *e*_𝑖,𝑗𝑗_ in PyMC (see code) which encodes the response arising from the 𝑖^𝑡ℎ^ electrode in the 𝑗𝑗^𝑡ℎ^subject. Within and between subject variances were quantified through hyper prior distributions placed on prior parameters^179^. Within and between subject variances were quantified through hyper prior distributions placed on prior parameters^179^.

#### 2.3 Prior selection

While LFPs elicited via INS, particularly in the auditory thalamocortical circuit, have largely been underexplored, previous knowledge to form highly informative prior distributions are not available. However, our extensive experience in INS single unit responses^3^, biophysical properties of auditory thalamus^45,155,166,182,183^ and cortex^46,72,183^, and recordings from and modeling of the central auditory pathway as a whole^163,184–186^ allow for the quantification of nominal thalamocortical firing properties into the prior distribution. ^45,155,166,182,183^ and cortex^46,72,183^, and recordings from and modeling of the central auditory pathway as a whole^163,184–186^ allow for the quantification of nominal thalamocortical firing properties into the prior distribution. As such, we have chosen to utilize moderately informative normally distributed prior distributions, allowing for the observed data to drive posterior inference. An alternative for noninformative prior distributions is the unform prior, which weighs all parameter outcomes as equally likely within the uniform interval. Normally distributed priors were chosen over uniformly distributed priors as all probabilities outside of the uniform interval are cast to 0 and are not included in the posterior distribution. Normal distributions put small, but not 0 probability on extreme events, which, if given enough evidence in the data, are represented in the posterior distribution. Initial observations of data showed that log transformation of predictor and response variables provided data distributions that were sufficiently normal, further facilitating the use of normally distributed priors. Prior sensitivity analyses also confirmed normally distributed priors did not have undue influence on posterior estimates.

#### 2.4 Posterior Decision Rules

Inference was conducted on posterior distributions utilizing 95% highest density interval estimation as measured through the maximum *a posteriori* (MAP) estimation standard to Bayesian practice^90,92^. Inference was conducted on posterior distributions utilizing 95% highest density interval estimation as measured through the maximum *a posteriori* (MAP) estimation standard to Bayesian practice^90,92^. Credible intervals, analogous to confidence intervals in frequentist inference, were measured as the highest density interval (HDI) containing 95% of the posterior distribution probability. A regression parameter was considered significant if the 95% HDI did not contain a value of 0, with zero indicative of a predictor having a null effect. MAP estimates, representing the mode of the posterior distribution and thus the most probable value of the regression parameter. This set of inference allows for the quantification of the uncertainty of parameter estimates, with narrow HDIs describing a more certain estimate of the regression parameter given the observed data.

#### 2.5 Final Statistical Models

Posterior predictive checking was performed by sampling the posterior distribution (20,000 draws) to create surrogate data distribution. Kernel density estimates of these posterior predictive distributions were compared to kernel density estimates of the observed data. Goodness of fit between the two distributions was assessed using Bayesian p-values quantifying the probability that posterior predictive-based draws are more extreme than observed data^91^. Posterior sensitivity to model prior and hyper prior distributions was assessed using leave-one-out cross (LOO) validation^94^. Models of standard regression with a normally distributed likelihood function and models of robust regression with Student-T distributed likelihood functions were compared with varying hyper prior variances. Robust regressions tend to perform better inference with less sensitivity to outlier data. Goodness of fit between the two distributions was assessed using Bayesian p-values quantifying the probability that posterior predictive-based draws are more extreme than observed data^91^. Posterior sensitivity to model prior and hyper prior distributions was assessed using leave-one-out cross (LOO) validation^94^. Models of standard regression with a normally distributed likelihood function and models of robust regression with Student-T distributed likelihood functions were compared with varying hyper prior variances. Robust regressions tend to perform better inference with less sensitivity to outlier data. Prior distributions for scale parameters of error and Student T tail mass 𝜈𝜈 were set to half-Cauchy distributions as Expected log pointwise predictive densities (ELPD)^187^ were calculated for each model.^187^ were calculated for each model. Higher values of ELPD are a result of higher out of sample predictive fit indicative of a better model. Weight values generated by LOO cross validation were also analyzed and predict the probability of each model given observed data. Finally, we observed the standard error of the ELPD estimate (SE), and the difference between the model with highest ELPD and every other model (dSE) with dSE of the top model set to 0.00 by definition. All LOO calculations were performed post hoc with the python package arviz, a plugin for PyMC. All models and hyperpriors are given in supplemental table I. Schematic descriptions of Bayesian hierarchical models is given in Figure S2. Posterior predictive Bayesian p-values for models are given in Figure S3,S4 respectively.

---

**Figure S1:**
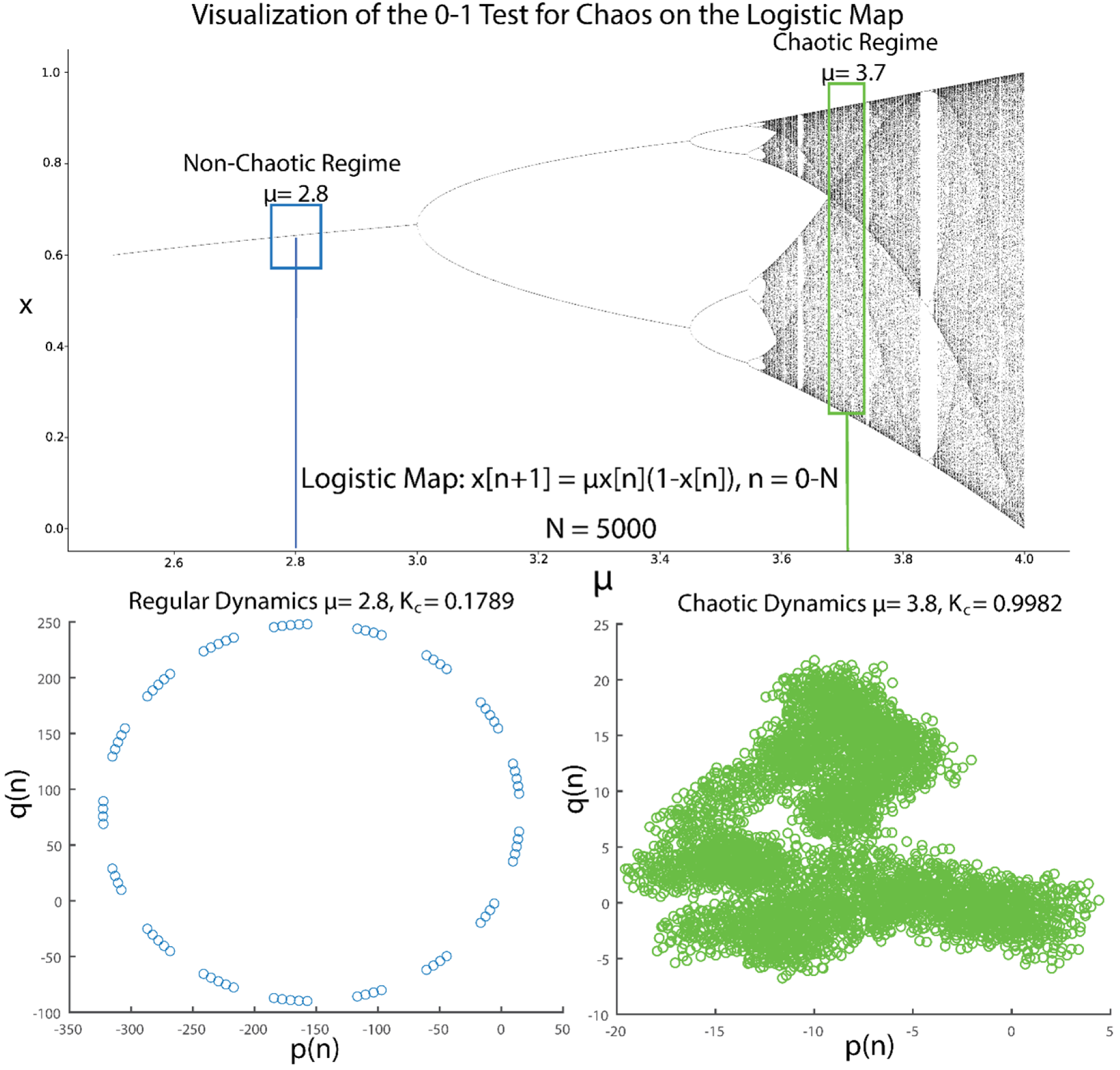
Visualization of the 0-1 Chaos test using the Logistic Map. The logistic map contains regions of regular, non-periodic and chaotic dynamics(top). Visualization of the logistic map cast into the 2D mapping p(n), q(n) shows that regular regimes show bounded dynamics with no growth in average mean-squared displacement with a K statistic close to 0 (0.1789)(bottom, left). Alternatively, chaotic regions of the logistic map cast into the p(n),q(n) space show dynamics with Brownian-like motion, and a linear growth in average mean-squared displacement, indicative of chaotic dynamics exhibiting diffusive growth of 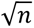. This is quantified with a K-statistic close to 1 (0.9982)(bottom, right).

**Figure S2:**
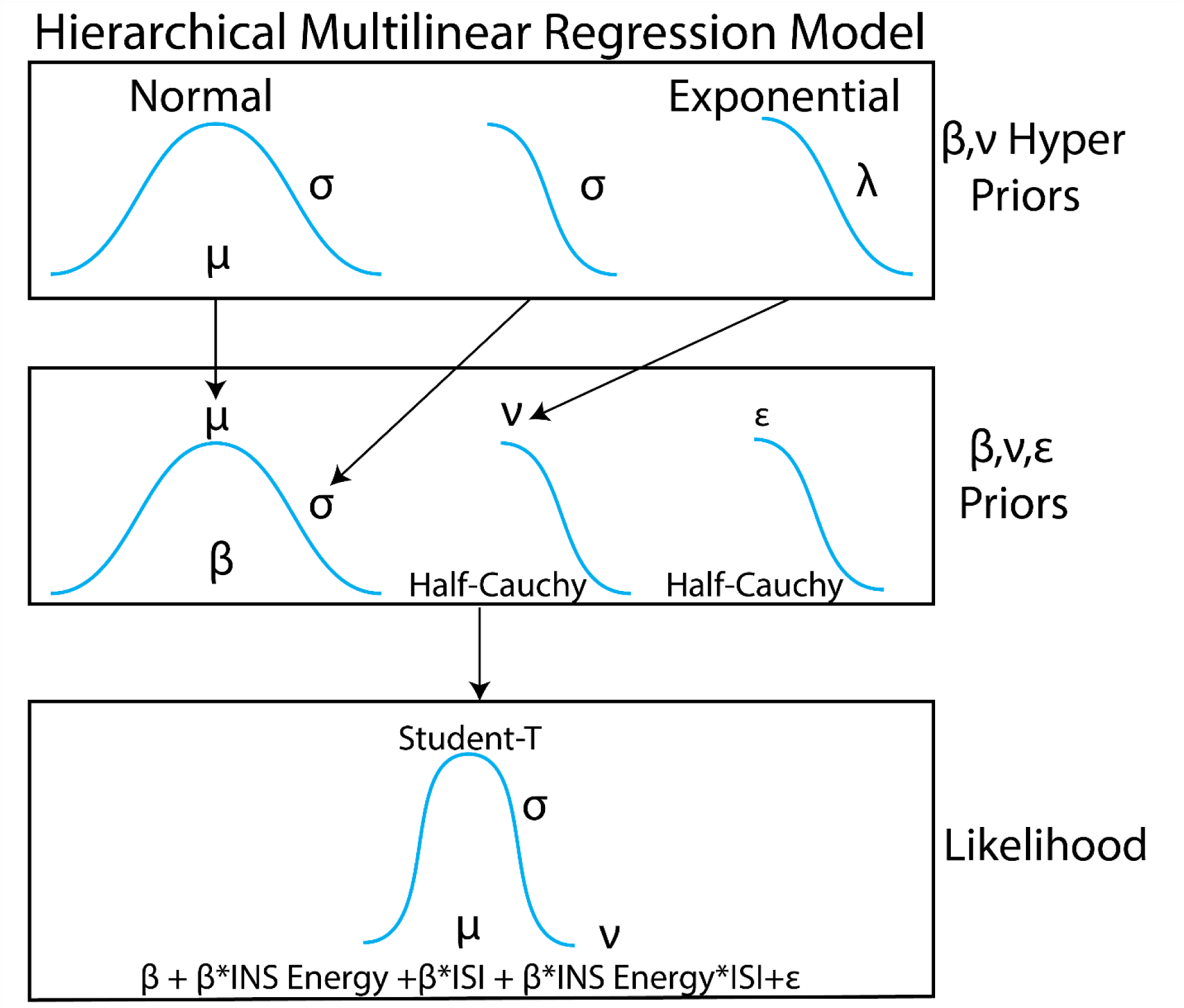
Schematic of Bayesian hierarchical linear regressions.

**Figure S3:**
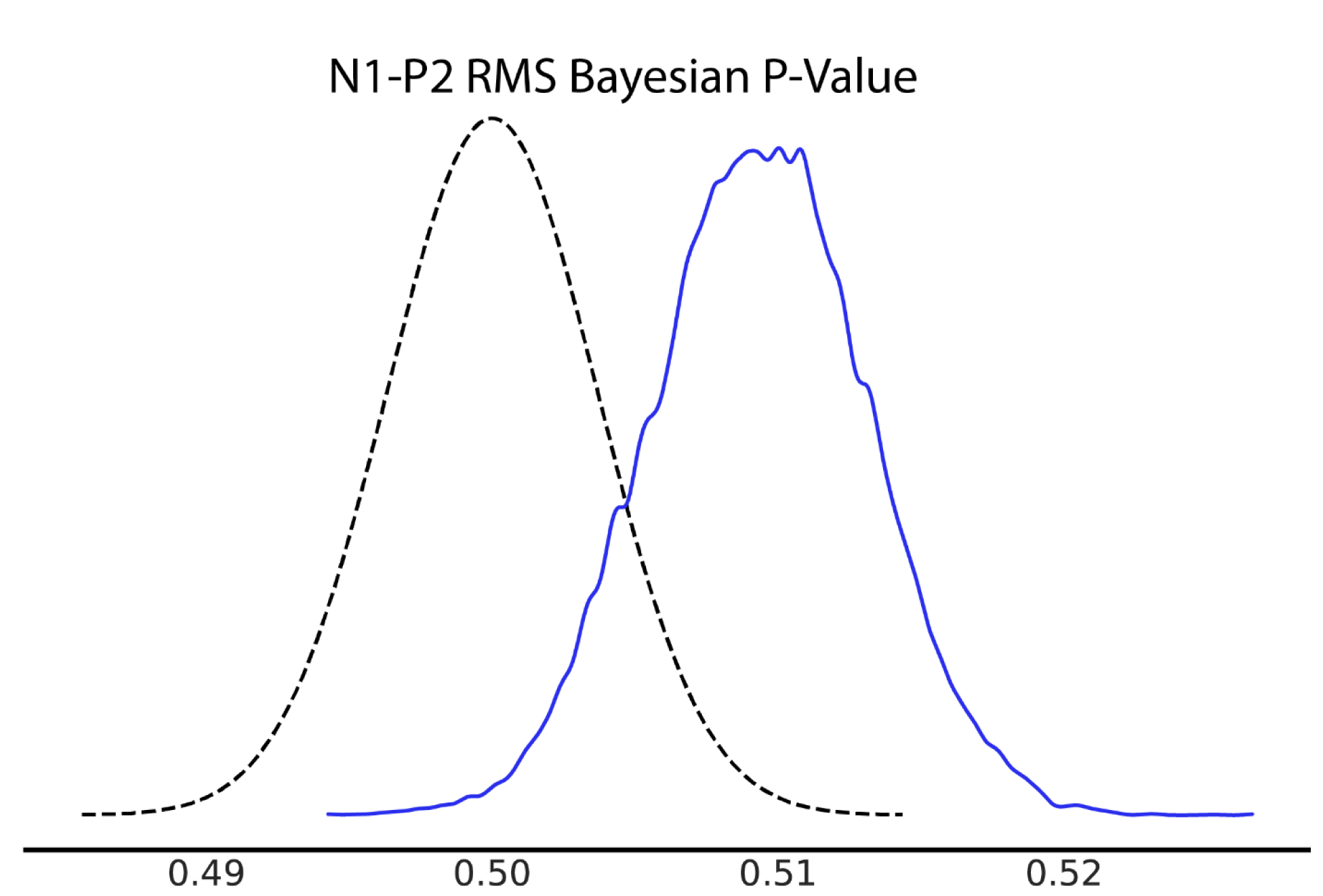
Bayesian p-value plot for N2-P1 LFP Bayesian hierarchical linear regression. Bayesian p-values suggest good model fit to data if posterior sampled distributions are near or centered about 0.5.

**Figure S4:**
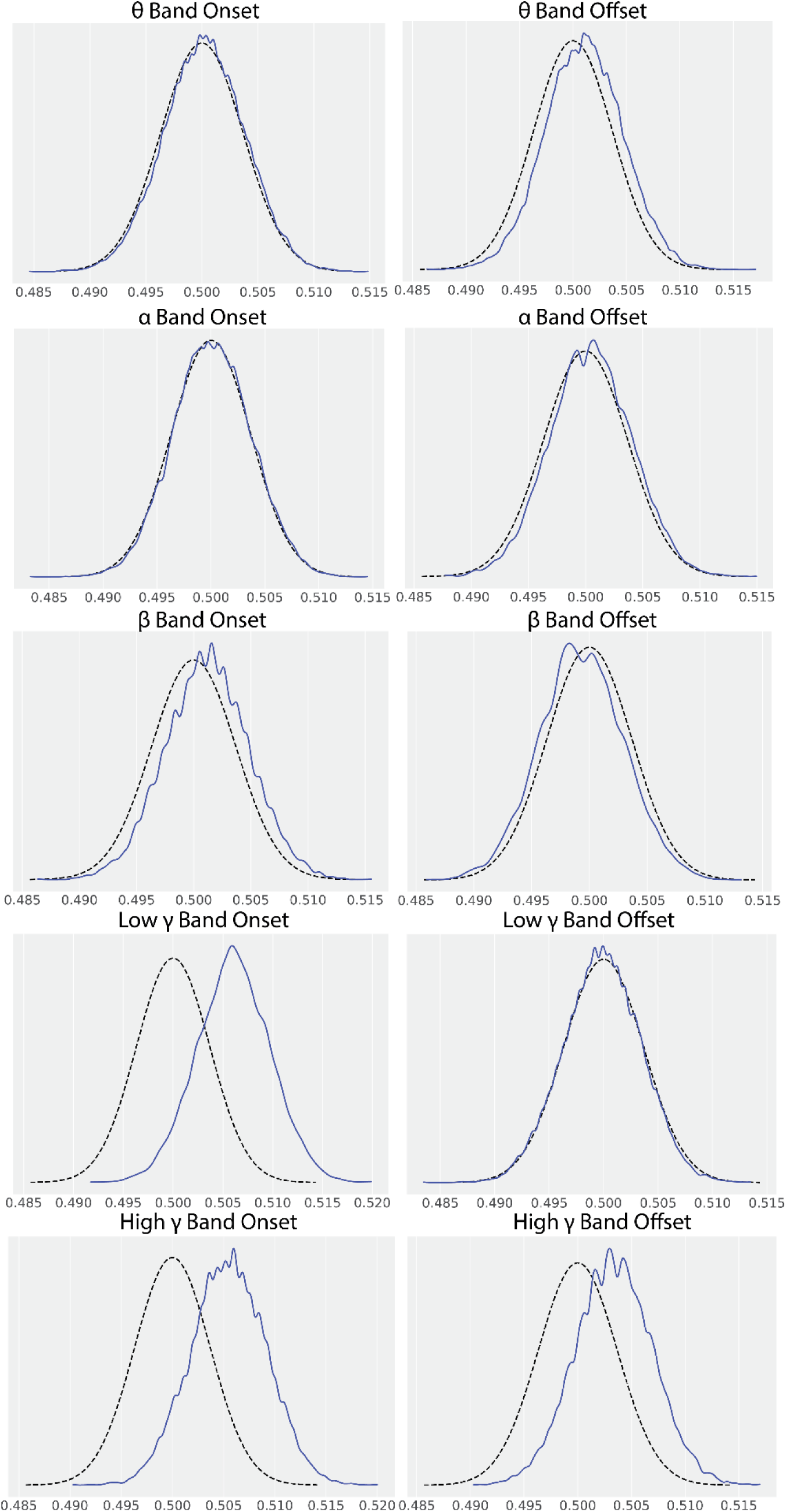
Bayesian p-values for LFP frequency band onset and offset statistical models.

**Figure S5:**
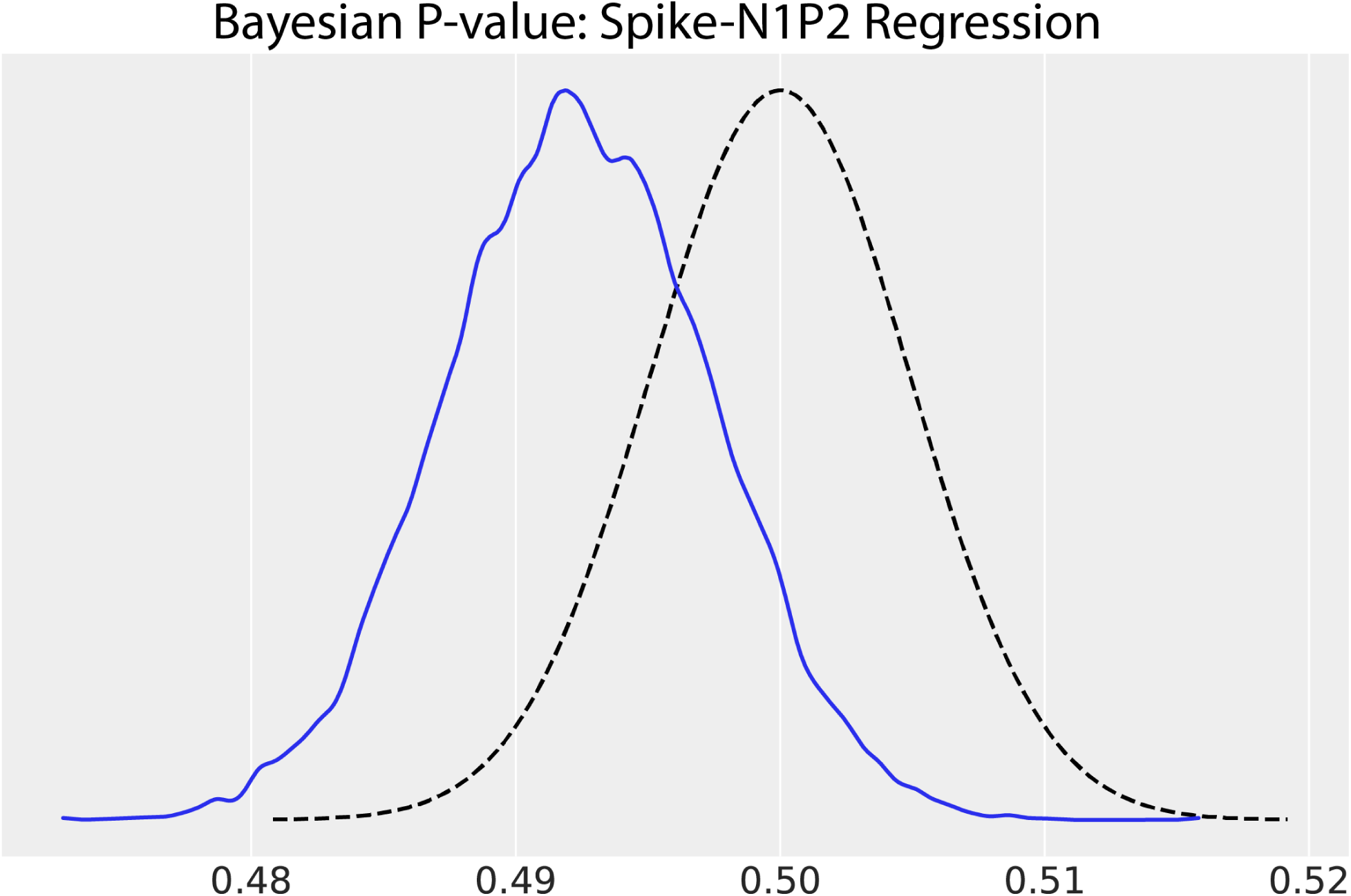
Bayesian p-values for spike rate-LFP N1-P2 RMS amplitude regressions.

**Figure S6:**
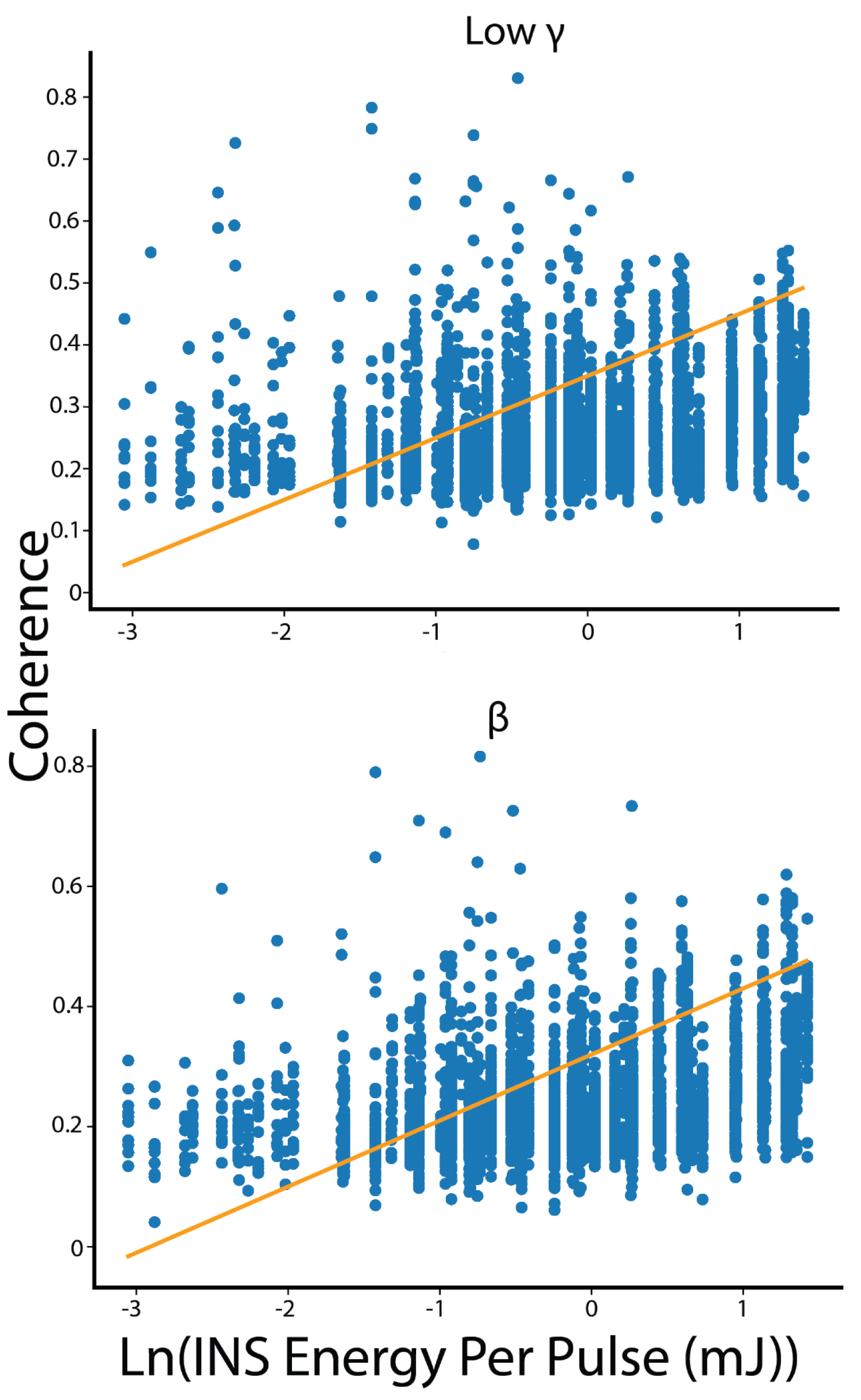
Hierarchical regression models for *β* and low *γ* spike-field coherences show evidence of nonlinearity as seen by low energy regression estimates which drop below measured coherences.

**Supplemental Table 1:** Final models used in statistical analyses. *ST*-Student T distributed, *N-*normally distributed.

| *Model Name* | *Likelihood* | *Final*<br>*Hyperpriors* |
| --- | --- | --- |
| *LFP N1-P2 dose response*<br>*curves* | $ST(\beta_{1,i}E + \beta_{2,i}ISI + \beta_{3,i}E * ISI + \varepsilon)$ | N(0,1) |
| **θ**-band LFP Onset | $ST(\beta_{1,i}E + \beta_{2,i}ISI + \beta_{3,i}E * ISI + \varepsilon)$ | N(0,10) |
| **θ**-band LFP Offset | $ST(\beta_{1,i}E + \beta_{2,i}ISI + \beta_{3,i}E * ISI + \varepsilon)$ | N(0,1) |
| **α**-band LFP Onset | $ST(\beta_{1,i}E + \beta_{2,i}ISI + \beta_{3,i}E * ISI + \varepsilon)$ | N(0,5) |
| **α**-band LFP Offset | $ST(\beta_{1,i}E + \beta_{2,i}ISI + \beta_{3,i}E * ISI + \varepsilon)$ | N(0,5) |
| **β**-band LFP Onset | $ST(\beta_{1,i}E + \beta_{2,i}ISI + \beta_{3,i}E * ISI + \varepsilon)$ | N(0,1) |
| **β**-band LFP Offset | $ST(\beta_{1,i}E + \beta_{2,i}ISI + \beta_{3,i}E * ISI + \varepsilon)$ | N(0,10) |
| Low **γ**-band LFP Onset | $ST(\beta_{1,i}E + \beta_{2,i}ISI + \beta_{3,i}E * ISI + \varepsilon)$ | N(0,1) |
| Low **γ**-band LFP Offset | $ST(\beta_{1,i}E + \beta_{2,i}ISI + \beta_{3,i}E * ISI + \varepsilon)$ | N(0,5) |
| High **γ**-band LFP Onset | $ST(\beta_{1,i}E + \beta_{2,i}ISI + \beta_{3,i}E * ISI + \varepsilon)$ | N(0,5) |
| High **γ**-band LFP Offset | $ST(\beta_{1,i}E + \beta_{2,i}ISI + \beta_{3,i}E * ISI + \varepsilon)$ | N(0,1) |
| Spike-field correlation<br>regressions | $ST(\beta_1 FiringRate + \beta_2 ISI + \varepsilon)$ | N(mean(firing<br>rate), 1) |
| SFC Spline Regression | $ST(\beta_i * w_i)$ | N(0,0.5) |

